# Membrane curvature catalyzes actin nucleation through nano-scale condensation of N-WASP-FBP17

**DOI:** 10.1101/2024.04.25.591054

**Authors:** Kexin Zhu, Xiangfu Guo, Aravind Chandrasekaran, Xinwen Miao, Padmini Rangamani, Wenting Zhao, Yansong Miao

## Abstract

Actin remodeling is spatiotemporally regulated by surface topographical cues on the membrane for signaling across diverse biological processes. Yet, the mechanism dynamic membrane curvature prompts quick actin cytoskeletal changes in signaling remain elusive. Leveraging the precision of nanolithography to control membrane curvature, we reconstructed catalytic reactions from the detection of nano-scale curvature by sensing molecules to the initiation of actin polymerization, which is challenging to study quantitatively in living cells. We show that this process occurs via topographical signal-triggered condensation and activation of the actin nucleation-promoting factor (NPF), Neuronal Wiskott-Aldrich Syndrome protein (N-WASP), which is orchestrated by curvature-sensing BAR-domain protein FBP17. Such N-WASP activation is fine-tuned by optimizing FBP17 to N-WASP stoichiometry over different curvature radii, allowing a curvature-guided macromolecular assembly pattern for polymerizing actin network locally. Our findings shed light on the intricate relationship between changes in curvature and actin remodeling via spatiotemporal regulation of NPF/BAR complex condensation.

## Introduction

Dynamic membrane curvature underpins vital cellular functions, orchestrating complex biophysical and biochemical activities ^1–4^. Membrane shape dynamics are intricately connected to the quick remodeling of the actin cytoskeleton, offering spatial and temporal governance of associated biological functions ^5–12^. Proteins that are sensitive to membrane curvature respond dynamically to nano-deformations, setting off intricate lipid-protein interactions at these sites. The mechanisms by which signal-triggered biomolecular cascades are locally intensified on the membrane and influence actin remodeling remain elusive.

BAR (Bin/Amphiphysin/Rvs) domain-containing superfamily of proteins provides an attractive template for studying the interplay between membrane curvature and the curvature-sensitive biochemical reactions. BAR domain proteins are known for their curvature-sensing and -shaping capabilities in cellular membranes, forming crescent dimers with lipid-affinity ^13–15^. These proteins, including subfamilies like BAR, N-BAR, F-BAR, and I-BAR, are central to processes such as endocytosis, membrane trafficking, and cell division ^16–19^. F-BAR proteins such as TOCA-1 and FBP17 and other BAR proteins including PACSIN, SNX9, amphiphysin, endophilin and IRSp53 were shown to interact with actin nucleator, and nucleation promoting factors (NPFs) and affect with actin polymerization ^20–27^. In addition, actin cytoskeleton-generated forces are also known to locally facilitate membrane morphogenesis ^7,21,28–33^. The direct role of membrane sculpting in triggering actin remodeling or perpetuating actin assembly around curved regions remains unclear, as decoupling the processes of curvature formation and actin polymerization is challenging.

The development of recent nanotopography technologies including nanopillars, nanobars, nanoridges, and nanobeads, each with a diameter of less than 500 nm, has allowed for exquisite control of substrate curvature ^6,9,12,34^. Such curvature control has demonstrated the selective recruitment of Wiskott-Aldrich syndrome protein (WASP) family proteins as a function of curvature ^5,6,11,12^. N-WASP is a membrane-bound actin nucleation-promoting factor, exists in an autoinhibited state ^35^. Activation occurs when it undergoes a conformational shift induced by its interaction with the small GTPase Cdc42 ^36^. Recent evidence suggests that N-WASP activation may also stem from multicomponent condensation, which was facilitated by influenced by its intrinsically disordered region (IDR)-mediated multivalent interactions. ^37–40^. But the role of multivalent interactions in controlling downstream actin signaling is poorly understood.

This study deciphered the spatiotemporal regulation of actin network formation at curved membrane sites with specific range of nanoscale radius. We reconstituted the curvature-dependent actin polymerization on a supported lipid bilayer integrated with precisely designed nanoarrays created using electron-beam lithography. Using mathematical modeling and quantitative biochemical and biophysical analysis, we discovered the step-by-step cascade reactions leading to curvature-mediated actin polymerization. Our findings show that this process involves precise curvature sensing by FBP17, the macromolecular condensation of the FBP17/N-WASP complex with an optimal stoichiometry in curvature-dependent manner, and the subsequent condensation-mediated activation of N-WASP for onsite actin nucleation. Our study reveals a novel mechanism in which nanoscale curvature precisely directs protein recruitment and macromolecular condensation to trigger actin nucleation locally. This work uncovers the intricate, dynamic interaction between membrane curvature and the biochemical cascade of actin polymerization that is fundamental to various cellular functions.

## Results

### The curvature radius-dependent recruitment of N-WASP requires FBP17

First, we built a minimal membrane curvature sensing system to investigate the molecular interactions between the curvature sensing of FBP17 and N-WASP. We used recombinant proteins of Arp2/3 NPF N-WASP (unless specifically stated, all the N-WASP used in our study was in an auto-inhibited state) and F-BAR family FBP17 and examined them over a defined gradient of membrane curvatures on nanobar arrays (Figure 1A,B). Such nanobars were fabricated through electron-beam lithography on quartz chips, containing two semi-cylindrical ends and a straight center. The radius of the curved ends was precisely controlled from 100 nm to 500 nm and the nanobars were 600 nm high. Similar nanoarrays demonstrated the recruitment of curvature-sensing proteins seen both in living cell systems ^6,9,41^ and *in vitro* on supported lipid bilayer (SLB) ^8^. Here, we prepared a supported lipid bilayer (SLB) on nanobar arrays similarly, featuring radius-gradient membrane curvatures on the ends of nanobars ^42^. We ensured that the lipid bilayer on specific curvature gradients maintained high fluidity, evidenced by a rapid fluorescence recovery after photobleaching (FRAP) (Supplementary Figure S1A,B). We then introduced Alexa Flour dye-labeled FBP17 (AF647-FBP17) and N-WASP (AF488-N-WASP) onto the lipid bilayer (Supplementary Figure S1C). To quantify curvature sensitivity, we averaged the fluorescent intensity of AF488-N-WASP and AF647-FBP17 over 50 nanobars and normalized it by the lipid bilayer signal to account for surface area differences. While the mean SLB signal was evenly distributed on the surfaces of nanobar, FBP17 and N-WASP exhibited an ends-enriched distribution dependent on the curvature radius (Figure 1C). The end density of FBP17 signal increases on sharper nanobars when the size is less than 500 nm (Figure 1D), aligning with previously reported curvature sensing ranges for FBP17 ^6,8^. Interestingly, N-WASP exhibited an even more prominent curvature response than FBP17 in high curvature regions less than 500 nm (Figure 1D,E), a range where FBP17 is known to sense ^8^. While, on the contrary, N-WASP alone does not exhibit curvature-dependent recruitment (Figure 1F,G).

**Figure 1.**
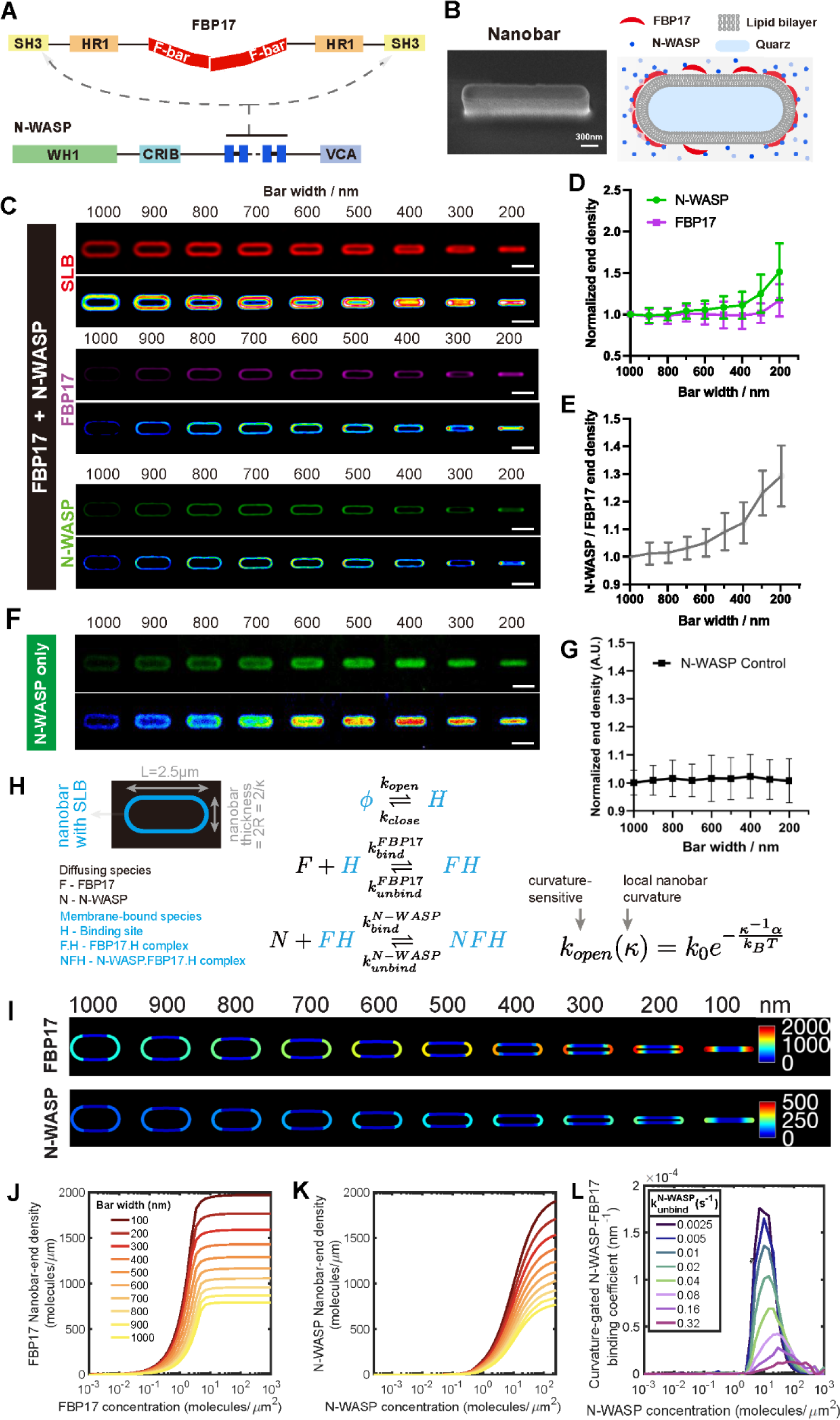
Understanding the curvature sorting of FBP17 and N-WASP through nano- patterning and spatial simulations. A. A domain diagram of FBP17 and N-WASP. The arrows indicate the interaction between SH3 domain of the dimeric FBP17 and the polyproline region of N-WASP. B. Left: Scanning electron microscopy image of a single nanobar. Tilted 30°. Scale bar: 300 nm. Right: A representation illustrating the *in vitro* reconstitution of FBP17-mediated N- WASP recruitment at membrane curvatures. C. Averaged confocal images showing curvature responses of SLB (upper), with additional 200 nM of AF647-FBP17, middle), or additional 50 nM of AF488-N-WASP (lower), respective on top of nanobars of 200-1000 nm width. Scale bar, 2 μm. D. Plot of normalized nanobar-end density of the FBP17 (purple) and N-WASP (green) in **(C)** as a function of nanobar width. Each point represents mean ± SEM from over 50 nanobars. E. Normalized N-WASP end density divided by FBP17 end density at nanobars of 200-1000 nm width in (D). Each point represents mean ± SEM from over 50 nanobars. F. Averaged confocal images showing curvature responses of 50 nM of AF488-N-WASP on top of nanobars of 200-1000 nm width. Scale bar, 2 μm. G. Plot of normalized nanobar-end density of N-WASP in **(F)** as a function of nanobar width. Each point represents mean ± SEM from over 50 nanobars. H. Spatial simulations considered two-dimensional SLB-coated nanobar (blue) with circular ends of radius R=bar width/2. FBP17 (F) and N-WASP (N) molecules diffuse in the free space around the nanbar and kinetically bind to the nanobar according to the following reactions. Curvature sorting is driven by curvature-sensitive exposure of hydrophobic patches (H) on SLB. FBP17 binds to H to form FH which then binds to N-WASP to form NFH. Please refer to Supplementary Methods for a detailed description of the model. I. Panel shows concentration profiles of FBP17 (top) and N-WASP (bottom) from simulations at various nanobar widths (mentioned on top). 𝑘_𝑏𝑖𝑖𝑛𝑑,𝐼_= 10 𝜇𝑚^2^/𝑚𝑜𝑙𝑒𝑐𝑢𝑙𝑒𝑠. 𝑠, 𝑘_𝑢𝑛𝑏𝑖𝑖𝑛𝑑,𝐼_= 0.01/𝑠, [𝐹𝐵𝑃17] = 1.1938 𝑚𝑜𝑙𝑐𝑢𝑙𝑒𝑠/𝜇𝑚^2^ J. Simulated profiles of nanobar-end density of FBP17 as a function of FBP17 concentration 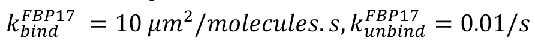 K. Simulated profiles of nanobar-end density of N-WASP as a function of N-WASP concentration ([FBP17]:[N-WASP] = 4:1). 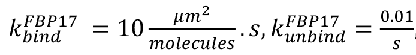 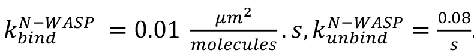 Please refer to Supplemental Figure S1I for profiles corresponding to other 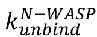 values. L. Concentration-dependent curvature-sensitivity is captured by calculating the binding coefficient obtained as follows. Given an N-WASP concentration, the ratio of FBP17 and N-WASP end-densities is plotted as a function of nanobar thickness. The plot shows profile of the slope at various N-WASP concentrations. **See also Supplementary Figure S1.**

To understand the relationship between the nanobar-induced curvature on the SLB and FBP17, we developed a computational model. To capture the curvature-sensitive nature of BAR domain proteins, without considering the structural and molecular domain details ^43–45^, we proposed a phenomenological curvature-sensitive binding reaction ^46^ with surface-volume coupling in two dimensions. To model the binding of FBP17 to curved SLBs, we consider the spontaneous appearance and disappearance of binding sites (H) on the membrane in a curvature-sensitive manner (Figure 1H). Freely diffusing FBP17 molecules in the volume can bind to the binding sites (H) on the SLBs and acts as substrates for N-WASP binding. These three reactions complete the VCell model ^47,48^. The geometries for the simulations were constructed similar to the experiment at FBP17:N-WASP=4:1 (Supplementary Figure S1D-H and Figure 1I). Our simulations revealed that indeed FBP17 density was higher on the curved regions of the nanopillar when compared to the flat regions and N-WASP followed the curvature-sensitive profile of FBP17 (Figure 1I). We next investigated how the density of FBP17 would vary on the curved regions as a function of FBP17 concentration and nanobar width. We found that that the curvature sensitivity of FBP17 was a function of FBP17 concentration and nanobar width (Figure 1J) and further predicted that N-WASP density followed a similar relationship (Figure 1K; Supplementary Figure S1I). To understand the [N-WASP]-dependent change to curvature sensitivity (𝐶 = ([𝑁 − W𝐴𝑆𝑃]/[𝐹𝐵𝑃17])_𝑛𝑎𝑛𝑜𝑏𝑎𝑟 𝑒𝑛𝑑_), we computed and plotted the slope of this curvature sensitivity,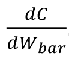, as a function of [N-WASP] and denoted this as a curvature-gated binding coefficient (Figure 1L). At low [N-WASP], very few FBP17 and N-WASP molecules are bound, showing no curvature sensitivity (𝑑𝐶/𝑑W_𝑏𝑎𝑟_∼0). Further, above a threshold [N-WASP], the nanobars are saturated with FBP17 and N-WASP molecules at all curvatures studied, resulting in curvature sensitivity inhibition, indicating a curvature-dependent stoichiometry of FBP17 to N-WASP. At intermediate concentrations, we see [N-WASP]-dependent curvature sensitivity (𝑑𝐶/𝑑W_𝑏𝑎𝑟_ > 0) and the [N-WASP] corresponding to peak curvature sensitivity depends on N-WASP unbinding rate (Figure 1L; Supplementary Figure S1I). Taken together, our simulations suggest that the curvature sensitivity of N-WASP emerges as a function of FBP17:N-WASP stoichiometric ratio and N-WASP kinetics.

### High membrane curvature promotes N-WASP binding by tuning (N-WASP:FBP17) bound stoichiometry

To validate the predictions from the model and to further investigate the interactions between FBP17, N-WASP and membrane curvature, we conducted experiments using a SLB with nano-array above. We systematically varied the concentrations of FBP17 and N-WASP while maintaining a constant molar ratio of 4:1 to investigate their radius-dependent recruitment as NPF on both patterned and flat SLBs (Figure 2A,B). The stoichiometry was selected based on a gradient titration of N-WASP, with concentrations ranging from 1 nM to 50 nM, and varying molar ratios to FBP17 (from 0.25 to 8) on flat SLBs (Supplementary Figure S2A). N-WASP showed a significant clustering at nano-scale levels at 50 nM and a molar ratio of F:N=4:1 (Supplementary Figure S2 B,C). Therefore, we chose the molar ratio of F:N=4:1 and adjusted N-WASP concentrations in the presence and absence of FBP17 on patterned SLBs and then analyzed the average intensity of the images obtained (Figure 2C-I). We quantified the density of FBP17 and N-WASP at the curved bar ends and normalized these measurements to lipid channel signals. By plotting the end density against protein concentration, we evaluated the accumulation of N-WASP mediated by FBP17 at the curved ends. As the curvature increased, both FBP17 (Figure 2D,E) and N-WASP (Figure 2F,G) exhibited greater accumulation at the curved ends when co-incubated on patterned SLBs. We observed that N-WASP accumulation is particularly sensitive to changes in curvature radius. This suggests a sequential recruitment mechanism, where N-WASP’s accumulation is contingent on FBP17’s primary detection of membrane curvature (Figure 2D,F). However, in the absence of FBP17, N-WASP no longer exhibited end-accumulation or curvature response (Figure 2H,I). In addition, we also engineered a gradient nano-pillar array, with diameters ranging from 200 nm to 1000 nm, to emulate FBP17-dependent N-WASP curvature sensing (Supplementary Figure S2D). Similarly, the N-WASP mirrored the curvature preference of FBP17, showing a ∼1.5-fold stronger accumulation on 200 nm pillars compared to the 1000 nm pillars (Supplementary Figure S2E,F). Together, we observed an increased N-WASP to FBP17 stoichiometry at high curvature areas of this membrane morphology (Supplementary Figure S2G), consistent with the bar-shape nano-patterning result.

**Figure 2.**
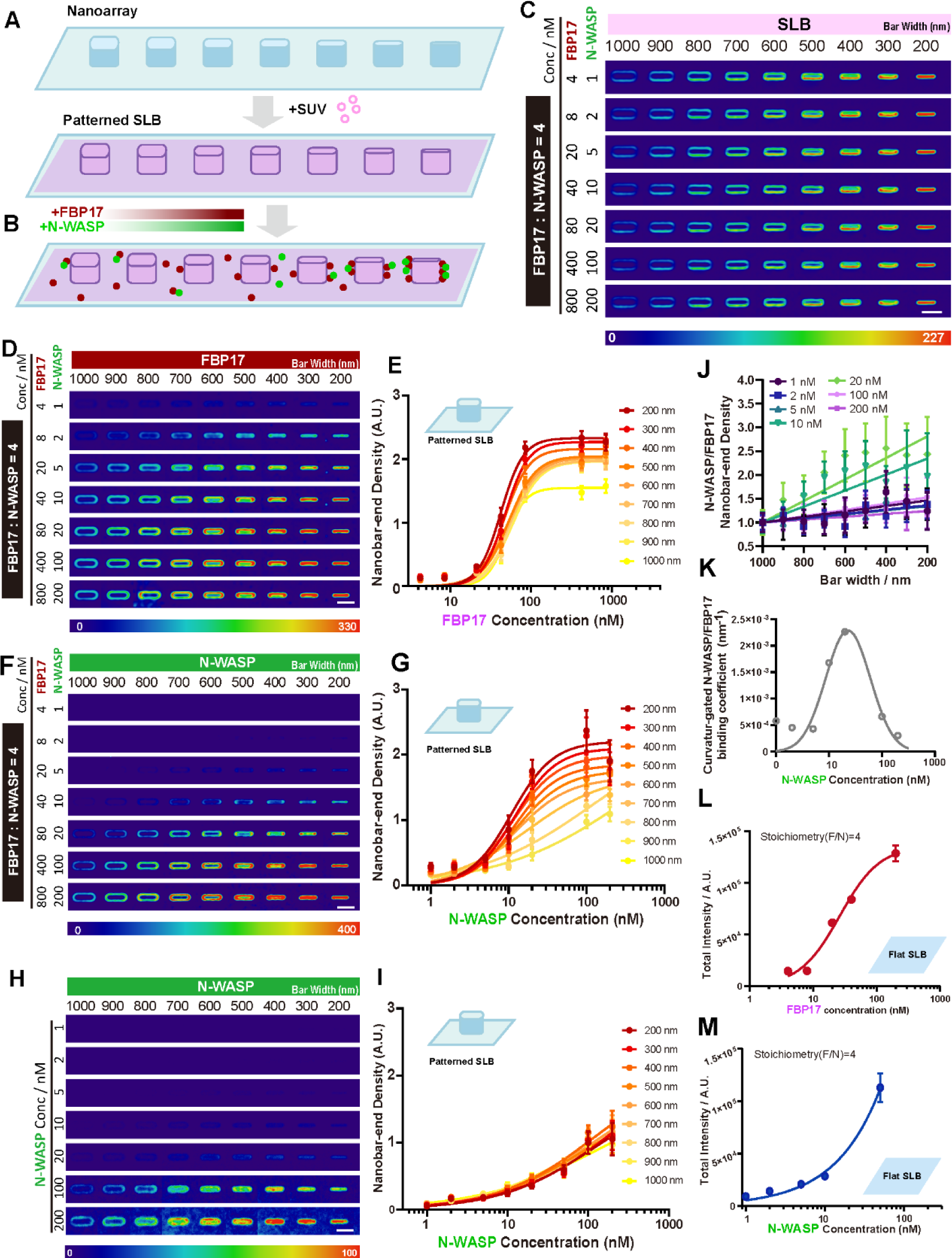
Curvature radius-dependent N-WASP and FBP17 distribution and condensation. A. Methodology illustration of patterned SLB B. Illustration of FBP17 and N-WASP titration on patterned SLB C. Averaged confocal images of lipid bilayer (10% POPS: 88.5% POPC: 1% PI(4,5)P2:0.5% Rhodamine-PE) on nanobars of 200-1000 nm in width. Scale bar, 2 μm. D. Averaged confocal images of 4, 8, 20, 40, 80, 400, 800 nM AF647-FBP17 (with 1, 2, 5, 10, 20, 100, 200 μM AF488-N-WASP under the molar ratio of F:N=4:1) on the lipid bilayer on top of nanobars of 200-1000 nm width. Scale bar, 2 μm. E. Plot of nanobar-end density of the 4, 8, 20, 40, 80, 400, 800 nM AF647-FBP17 (with 1, 2, 5, 10, 20, 100, 200 nM AF488-N-WASP under the molar ratio of F:N=4:1) as a function of concentration. Lines are binding curves fitted with the Hill equation. Each point represents mean ± SEM from over 50 nanobars. F. Averaged confocal images of 1, 2, 5, 10, 20, 100, 200 nM AF488-N-WASP (with 4, 8, 20, 40, 80, 400, 800 μM AF647-FBP17 under the molar ratio of F:N=4:1) on the lipid bilayer on top of nanobars of 200-1000 nm width. Scale bar, 2 μm. G. Plot of nanobar-end density of the 1, 2, 5, 10, 20, 100, 200 nM AF488-N-WASP (with 4, 8, 20, 40, 80, 400, 800 nM AF647-FBP17 under the molar ratio of F:N=4:1) as a function of concentration. Lines are binding curves fitted with the Hill equation. Each point represents mean ± SEM from over 50 nanobars. H. Averaged confocal images of 1, 2, 5, 10, 20, 100, 200 nM AF488-N-WASP control on the lipid bilayer on top of nanobars of 200-1000 nm width. Scale bar, 2 μm. I. Plot of nanobar-end density of the 1, 2, 5, 10, 20, 100, 200 nM AF488-N-WASP as a function of concentration. Lines are binding curves fitted with the Hill equation. Each point represents mean ± SEM from over 50 nanobars. J. Normalized N-WASP end density divided by FBP17 end density at nanobars of 200-1000 nm width under 1, 2, 5, 10, 20, 100, 200 nM N-WASP concentration with same stoichiometry N/F=1/4. K. Curvature responsive coefficient plot (slope of N-WASP/FBP17 ratio as a function of concentration). Lines are binding curves fitted with the Gaussian equation. Each point represents mean ± SEM from over 50 nanobars. L. Plot of single particle total intensity of 4, 8, 20, 40, 200 nM AF647-FBP17 (with AF488-N- WASP under the molar ratio of F:N=4:1) on flat lipid bilayer(10% POPS + 89.5% POPC + 0.5% Rhodamine-PE). Each point represents mean ± SEM, N=200 randomly selected from over 10,000 particles. M. Plot of single particle total intensity of 1, 2, 5, 10, 50 nM AF488-N-WASP (with AF647- FBP17 under the molar ratio of F:N=4:1) on flat lipid bilayer (10% POPS + 89.5% POPC + 0.5% Rhodamine-PE). Each point represents mean ± SEM, N=200 randomly selected from over 10,000 particles. **See also Supplementary Figure S2.**

We further explore how the progressive assembly of the macromolecular FBP17/N-WASP complex enhances the dynamic recruitment of N-WASP molecules in relation to membrane curvature, taking into account both curvature and protein concentration gradients. We measured the N/F ratio, which reflects the relative number of N-WASP molecules associated with each FBP17 molecule. Our results showed that the N/F ratio increases with membrane curvature under all the protein concentration conditions we examined (Figure 2J), confirming that curvature promotes N-WASP recruitment to FBP17. In line with our computational models, we found that the rate of increase in the N/F ratio with curvature—a marker of the molecular assembly of the clusters—is also influenced by the local concentration of the proteins (Figure 2J). This suggests that membrane curvature can fine-tune the stoichiometry between FBP17 and N-WASP molecules, reaching an optimized equilibrium. Consistent with the simulation results (Figure 1L), we identified a specific concentration range where FBP17 optimally facilitates N-WASP clustering. The point at which the N/F ratio’s response to curvature peaks is when N-WASP is at 20 nM in the presence of 80 nM FBP17 (Figure 2K). Above these concentrations, the responsiveness of the N/F ratio to curvature decreases, suggesting a potential geometric constraint limiting the assembly of the FBP17/N-WASP complex on curved membrane areas, over a broad range of nanomolar concentrations.

To investigate whether the constrained progression results from the 3D membrane geometry generated by curvature, we conducted measurements of N-WASP single particle intensity on a flat SLB. Within this 2D flat membrane system, N-WASP did not exhibit a saturation state and continued to accumulate over the same stoichiometry range with FBP17 (Figure 2L,M). The collective data indicate that curvature geometry is crucial not just for facilitating the macromolecular assembly of the FBP17-N-WASP complex but also for modulating the extent of their assembly. It further influences the interaction dynamics and conformation of FBP arrays encircling the curvature, thereby dictating their stoichiometry through topographical cues. In summary, N-WASP localizes prominently in response to positive curvature, forming percolation clusters that depend on the presence of FBP17, high membrane curvatures, and protein concentration.

### Actin polymerization in a radius-dependent manner

We next investigated the functional relationship between membrane curvature and dynamic actin remodeling in living cells. Previous studies have shown that F-actin in U2OS cells exhibits greater accumulation at high membrane curvatures ^6^. To dissect the curvature-guided initiation of dynamic actin polymerization and network organization, we first treated U2OS cells with Latrunculin A (Lat A) to depolymerize the F-actin and then monitor F-actin recovery by following the Lat A washout (Figure 3A). First, lifeact-mApple labeled F-actin bundles in cells on flat surfaces outside of curvature zones and also highlighted dense F-actin networks at the end of high-curvature regions above the nanobars (Supplementary Figure S3A, left panels). If we slowly depolymerize F-actin by treating cells at a non-toxic low-dose of LatA (100 nM), F-actin signals were no longer visible at the curved ends (Supplementary Figure S3A, middle panels,). Next, we allowed a repolymerization of actin network upon Lat A washout. We observed a pronounced F-actin recovery around the nano-pattern in a radius-dependent manner (Figure 3B-D, Supplementary Figure S3A, right panels), indicating an efficient onsite nucleation of actin cytoskeleton at the high curvature regions.

**Figure 3.**
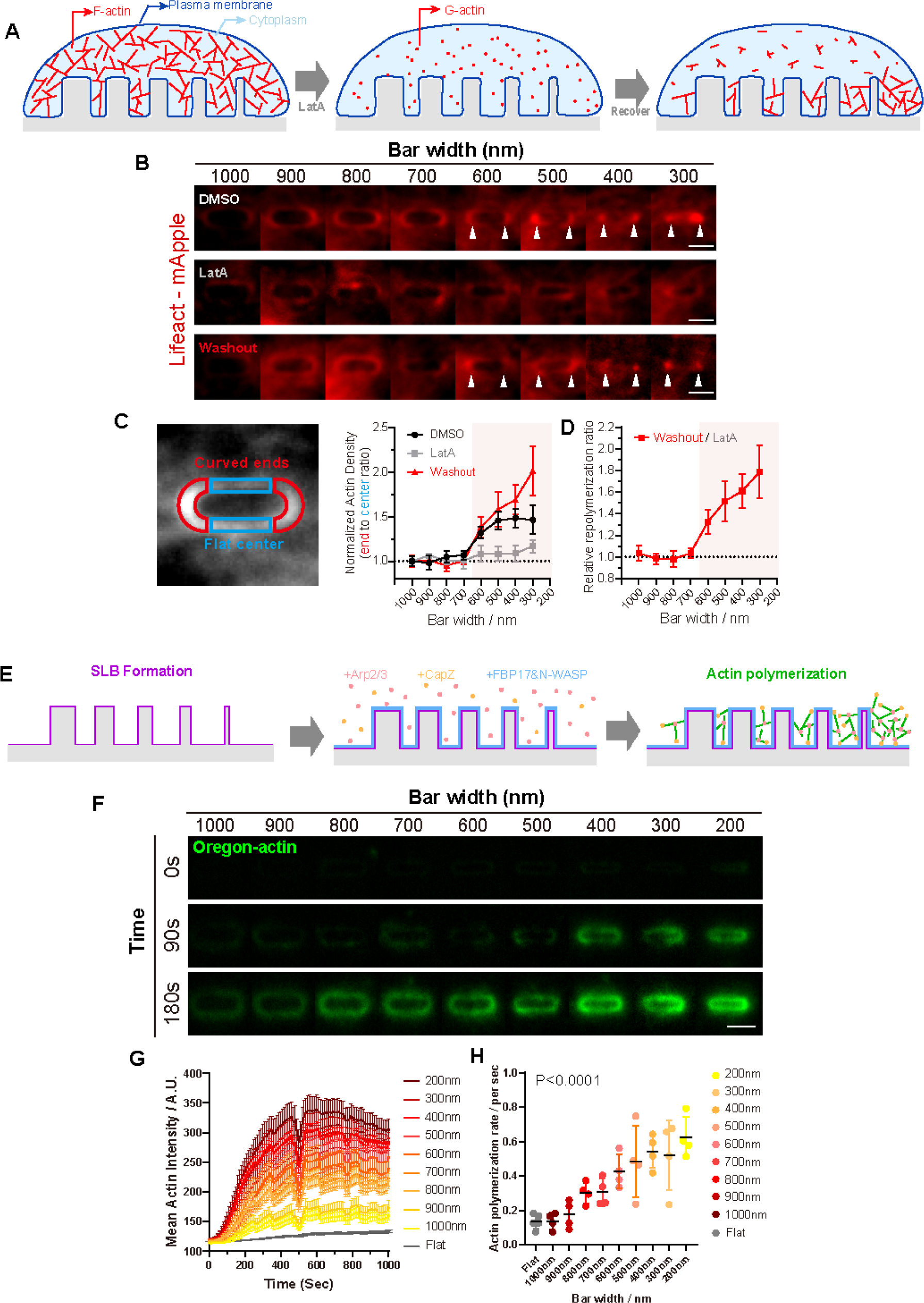
Radius-dependent actin assembly. A. A schematic illustrates the process of in vivo LatA treatment assay in U2OS cell lines on nanostructures. B. Averaged confocal images of LifeAct-mApple-expressed U2OS live cells cultured on nanobars ranging from 300 to 1000 nm in width demonstrate F-actin recovery following a 1h treatment with 100 nM LatA and a 20min washout. Scale bar, 2 μm. C. Normalized actin signal density calculated based on the actin intensity in the flat area, at nanobars ranging from 300-1000 nm in width. Each point represents mean ± SEM from over 50 nanobar ends. D. The normalized actin repolymerization ratio (Washout / LatA) in (C) at nanobars ranging from 300-1000 nm in width. Each point represents mean ± SEM from over 20 nanobar ends. E. A schematic illustrates the process of in vitro reconstitution of FBP17-mediated actin polymerization at membrane curvatures. F. Averaged confocal images of in vitro reconstitution of 1.5 μM actin (10% Oregon labeled) polymerization at 0/90/180s on the bilayer at nanobar of 200-1000 nm width with the presence of 200 nM FBP17, 50 nM N-WASP, 12 nM CapZ and 5 nM Arp2/3. Scale bar, 2 μm. See also Movie S1. G. Normalized actin signal intensity based on their corresponding lipid bilayer intensity at nanobars of 200-1000 nm width. Each point represents mean ± SEM from 4 nanobars. H. Actin polymerization rate at nanobars of 200-1000 nm width. (The linear fitted slope within 105-350s range in G). Each point represents mean ± SD, N=4. **See also Supplementary Figure S3.**

We were next motivated to dissect the detailed biochemical reactions underlying the above living cell results; we established an *in vitro* biochemical reconstitution assay for radius-dependent actin polymerization, enabling quantitative analysis of the curvature and FBP17-guided nucleation-activities of N-WASP. We created a SLB with a curvature- gradient nanobar array and applied G-actin and recombinant proteins, including FBP17, N-WASP, Arp2/3, and CapZ, to examine NPF activation for actin nucleation (Figure 3E). The presence of capping protein limited the initial elongation and spontaneous polymerization of F-actin, allowing a nucleation-specific assessment of its efficiency. Thus, we can monitor the initiation and accumulation of actin signals around nanobars, spanning from 200 nm to 1000 nm, over time and observe within 180s, the initiation period of actin polymerization (Figure 3F, Movie S1). We noticed that 200-400 nm nanobars showed rapid actin polymerization starting from 90s, while wider ones took longer, which indicates that actin nucleation is accelerated on highly curved membranes (Figure 3F). When actin signal was within initial linear increase period, curvature-dependent actin assembly was shown by quantifying polymerization rate (Figure 3G,H). In contrast, non-polymerizable(NP)-G-actin-based polymerization assay did not show obvious actin signal increase over curvature radius (Supplementary Figure S3B-D). We additionally compared the conditions without curvature and without capping proteins, respectively. On the flat SLB, we only observed spontaneous actin polymerization after 15min (Supplementary Figure S3E left), whereas an additional FBP17 exhibited randomly distributed actin asters, suggesting activated N-WASP in F-actin nucleation (Supplementary Figure S3E right). In the absence of CapZ, spontaneously polymerized F-actin distributed randomly and connected on the flat SLB, whereas on SLB-coated nanobar arrays, actin cytoskeleton is highly aligned with curvature end of nanobars (Supplementary Figure S3F). The above *in vivo* and *in vitro* reconstitution suggested a FBP17-mediated activation of N-WASP that is highly coupled with membrane curvature, which provides a hotspot for the reaction and also guides components stoichiometry with optimized activities on demand, in space and time.

### FBP17 amplifies N-WASP-mediated actin assembly by forming condensed nucleator complex zones

Given the demonstrated multivalent interactions for the FBP17/N-WASP complex (Figure 1A), along with the notably increased F-actin polymerization, which occurs in a curvature radius-dependent manner (Figure 3D, H), we are motivated to explore the specific molecular mechanisms that direct this cascade of recruitment and regulate the biochemical activity of actin polymerization (Figure 4A). Upon investigating the high curvature nanobar, we noted that FBP17 exhibited a ∼1.2-fold stronger signal end density at 200 nm in comparison to that observed at a nanobar size of 1000 nm. In the case of N-WASP, this ratio saw an increase to approximately 1.5-fold, and for the rate of actin polymerization, the ratio further rose to around 2.7-fold (Figure 4B). Thus, we hypothesize that FBP17- driven N-WASP localization is critical to amplifying actin nucleation.

**Figure 4.**
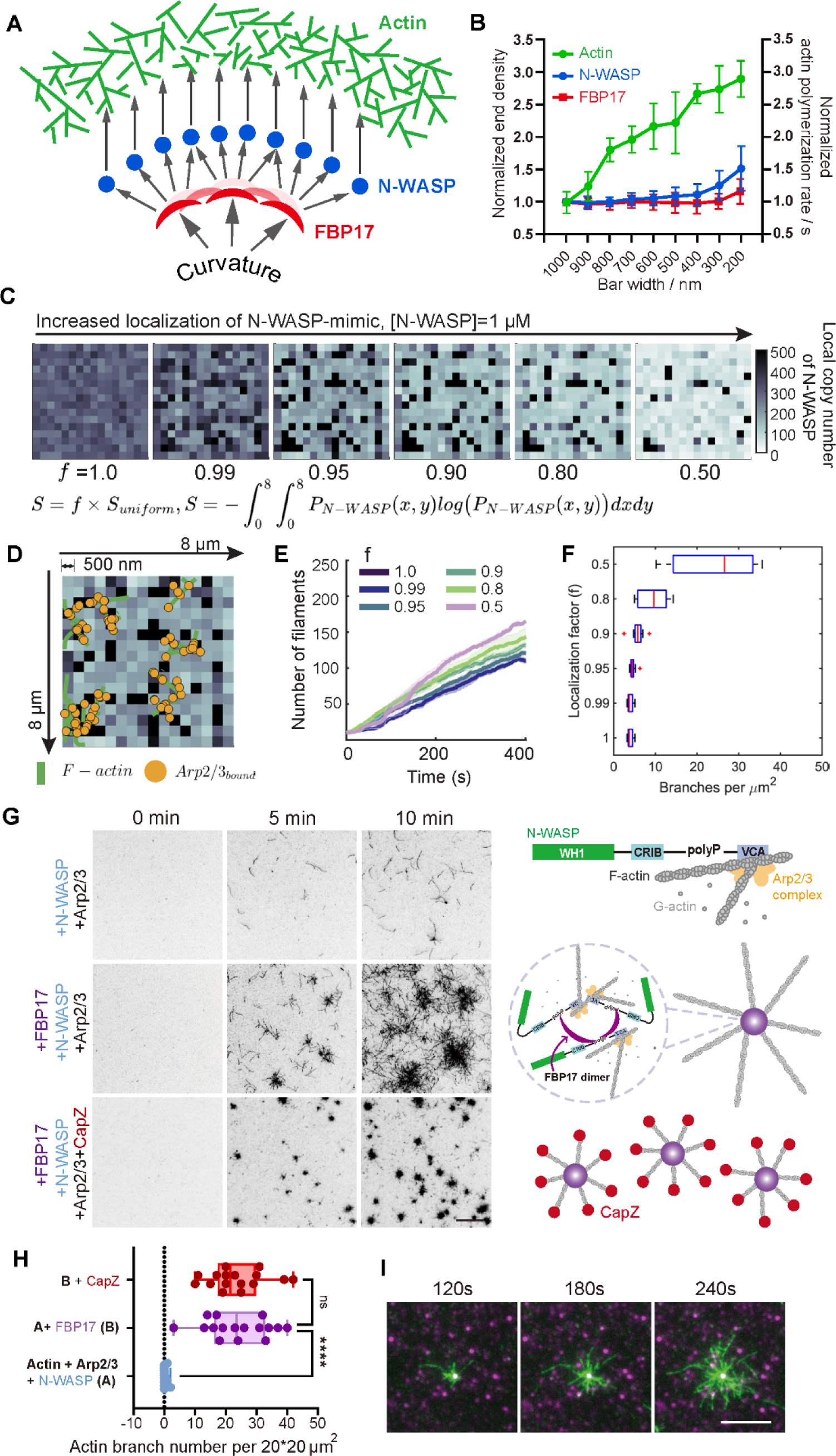
FBP17 localization activates N-WASP clustering resulting in enhanced actin nucleation. A. A schematic illustrates the in vitro reconstitution of amplification effects for actin polymerization at membrane curvatures. B. Normalized nanobar-end density of FBP17(red) and N-WASP(blue) based on their corresponding lipid bilayer intensity (each point represents mean ± SEM from over 50 nanobars) and normalized actin polymerization rate(green) at nanobars of 200-1000 nm width (each point represents mean ± SEM from 4 nanobars). C. Spatial clustering of N-WASP is mathematically represented using the SHannon entropy function. Spatial density maps of N-WASP-mimics were subsequently generated at various values of the spatial localization factor (f) using constrained optimization technique (Please refer to Supplementary Methods for detailed information). Colormaps show resulting N- WASP densities are shown for 8 μm x 8 μm space divided into compartments of size 0.5 μm x 0.5 μm. D. Representative final snapshot (t=400s) corresponding to MEDYAN simulation of actin chemical dynamics at f=0.9 in shown. The underlying N-WASP-mimic density is shown along with overlay of F-actin filaments (green) and filament-bound Arp2/3 (yellow). E. Mean and standard error of mean time series profiles of number of filaments are shown colored by spatial localization factor (f) (N=5). F. Box plot shows distribution of branches per cluster computed from simulations with varying spatial localization of N-WASP. Data shown is scaled for 20x20 μm^2^ cluster for ease of comparison against experiments. Red line represents median, the box bounds represent quartiles, and the whiskers represent bounds corresponding to 95% confidence intervals. Outliers are shown as red crosses. G. TIRF images of actin polymerization (10% Oregon-labeled) with different combinations of 80 nM FBP17, 20 nM N-WASP, 5 nM Arp2/3 and 3 nM CapZ. Scale bar, 10 μm. See also Movie S2. H. Distribution of number of branches (mother and attached branches) at t=300 s per 20*20um2. N=10, Mean ± SD shown. I. Zoom-in time-lapse TIRF images of actin clusters (10% Oregon-labeled) with AF647- FBP17 as a center. Scale bar, 5 μm. **See also Supplementary Figure S4.**

To test if N-WASP localization can enhance Arp2/3-driven nucleation, we generated 8 × 8 𝜇𝑚^2^ copy number maps of N-WASP that were spatially discretized into compartments of size 0.5 × 0.5 𝜇𝑚^2^for different levels of localization. Towards this, we quantify localization based on the Shannon Entropy, 𝑆(𝑥, 𝑦) of the corresponding probability density functions. Uniform distributions result in maximum value 𝑆_𝑚𝑎𝑥_ of the Shannon entropy function. We generated probability density functions corresponding to various levels of N-WASP-localization using least squares optimization of the objective function 𝐺: = (𝑆 − 𝑓𝑓 × 𝑆_𝑚𝑎𝑥_)^2^, 𝑓𝑓 ∈ [0,1]. Here, the factor *f* is termed as the localization factor and S represents the Shannon entropy of a given probability density. f=1 corresponds to an objective function whose minima corresponds to uniform distribution. Please refer to Supplementary Methods for a detailed description of the model. The resulting probability density maps were sampled to generate copy number maps corresponding to [𝑁 − W𝐴𝑆𝑃] = 1 𝜇𝑀 (Figure 4C). We then simulated the resulting Arp2/3 nucleation using MEDYAN (Mechanochemical Dynamics of Active Matter), a C++ code to study the stochastic mechano-chemistry of active networks (Figure 4D) ^49–51^. Initial condition for simulations at each localization factor consists of the N-WASP copy number map generated along with uniformly distributed G-actin ([𝐺 − 𝑎𝑐𝑡𝑎𝑎𝑛] = 1 𝜇𝑀) and inactive Arp2/3 ([𝐴𝑟𝑝2/3_𝑖𝑖𝑛𝑎𝑐𝑡𝑖𝑖𝑣𝑒_] = 10 𝑛𝑀) molecules. We consider N-WASP-dependent activation of Arp2/3 molecules along with polymerization, depolymerization, and dendritic nucleation reactions as described in Supplementary Figure 4A. Our simulations reveal that the number of filaments nucleated increases with localization of N-WASP-Arp2/3 (lower f value, Figure 4E). Additionally, we also show that there is a threshold activation rate of Arp2/3 below which the localization effect is inhibited (Supplementary Figure 4B-D). Finally, we identified the branch clusters in the reaction volume and calculated the number of branches per 𝜇𝑚^2^. Our findings show that the branch density also increases with increased localization (Figure 4F).

We also experimentally examine these model predictions by reconstituting an N-WASP-mediated Arp2/3 complex activation to study actin nucleation. We first sought to investigate if FBP17 could facilitate N-WASP-mediated activation of actin assembly. By experimenting with various mixtures of FBP17, N-WASP, and the Arp2/3 complex, we found that FBP17 triggers the formation of aster-shaped clusters of actin filaments (Figure 4G,H; Movie S2). Each cluster of actin filaments had an FBP17-enriched focus acting as the origin of nucleation, mirroring the accumulation pattern of FBP17 on a flat SLB (Figure 4I and Supplementary Figure S3F left). This observation suggests that FBP17 initiates actin polymerization by forming individual N-WASP-based nucleation centers through macromolecular assembly. Furthermore, we noted that the aster-shaped actin clusters would still form even in the presence of CapZ (Figure 4G). CapZ binds to the actin barbed end and hinders the elongation of actin seeds outside of the FBP17/N-WASP nucleation center, around which the filaments are also shorter (Figure 4G). Additionally, we also see that the increase in the number of branches in the presence of FBP17 driven N-WASP clusters (Figure 4G-H) is consistent with predictions from our model (Figure 4C-E).

### N-WASP condensation is mediated by multivalent interaction with FBP17

To investigate the cascade condensation FBP17 and N-WASP complex for nucleating Arp2/3-mediated actin polymerization, we next examined whether homo-dimeric FBP17 can cluster full-length N-WASP through the direct interaction between its SH3 domains and N-WASP disordered PolyP region (271-391aa) (Figure 1A). Using TIRFM imaging on SLB with the same lipid components as above, the size and total intensity of N-WASP foci increased with the addition of nanomolar FBP17 after 15min incubation (Figure 5A,B). Although the N-WASP signal is partially colocalized with FBP17, their colocalization generates bigger and more condensed foci (Figure 5C,D). Meanwhile, we found FBP17 also exhibits clustering behavior after adding N-WASP, and the size of FBP17 clusters is grown stoichiometric (Supplementary Figure S5A,B). This observation revealed that FBP17 can induce the condensation of N-WASP molecules, which could be a potential mechanism for FBP17-mediated activation of N-WASP.

**Figure 5.**
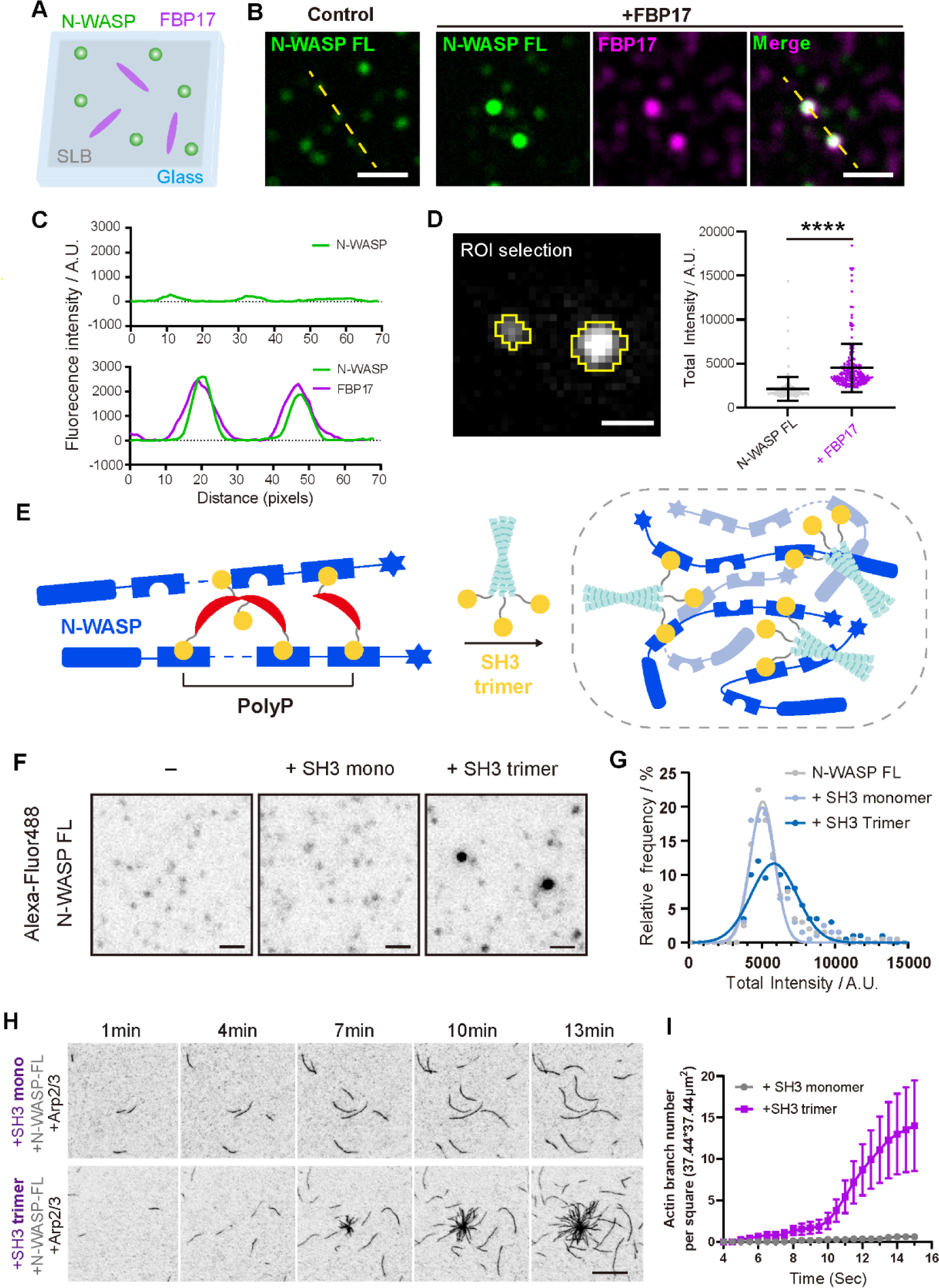
N-WASP condensates mediated by FBP17 multivalent interaction. A. A schematic illustration of protein assembly assay on flat SLB. B. Single particle TIRF images of 5 nM AF488-N-WASP with 20 nM AF647-FBP17 on SLB. Scale bar, 2 μm. C. Intensity profiles along the dashed yellow lines shown in (B) showing that clustering of N- WASP occurred in the presence of FBP17 and N-WASP colocalized with FBP17 in the clusters. D. Left: the Roi selection of single particle by threshold. Right: total intensity plot of N-WASP single particles on SLB. N=200 out of over 10,000 particles from 2 repeats, SD showed. E. A schematic illustrates the multivalent interaction between N-WASP (blue) and FBP17 (red) or SH3 trimer (yellow). F. Single particle TIRF images of 5 nM AF488-N-WASP with 20 nM AF647-SH3 or 20 nM AF647-oTri-SH3 on SLB. Scale bar, 1 μm. G. Distribution of N-WASP single particles population on SLB. N=200 out of over 5000 particles from 2 repeats, gaussian normalization applied. H. TIRF images of actin polymerization (10% Oregon-labeled) with different combinations of 80 nM SH3 or oTri-SH3, 20 nM N-WASP and 5 nM Arp2/3. Scale bar, 10 μm. See also Movie S3. I. Number of filaments at t=60-960s per 37.44*37.44um^2^. N=10, Mean ± SEM shown. **See also Supplementary Figure S5 and S6.**

However, FBP17-mediated clustering is theoretically not the only factor that can trigger N-WASP activation. To differentiate the mechanisms of FBP17-mediated-clustering and known conformational exposure in activating N-WASP, we generated additional three truncating variants, including auto-activated N-WASP L235P, PI(4,5)P2-binding-activated CRIB-PolyP-VCA (N-WASP 151-505), and verprolin-cofilin-acidic (VCA) ^36,52^ (Supplementary Figure S5A). To examine whether the conformational and lipid-binding activation mechanisms mediate the observed N-WASP clustering, we titrated the N-WASP WT and truncations on flat SLB with a same range of concentration. Our experiments revealed that in contrast to the FBP17-mediated condensation of N-WASP WT, neither N-WASP WT nor its truncated variants lacking the conformational and lipid-binding activation mechanisms managed to form higher-order assemblies (Supplementary Figure S5C). These findings underscore the distinctive N-WASP activation mode via FBP17-guided macromolecular assembly. We next asked whether the multivalent interaction of SH3 and PolyP plays a determinative role in this nano-condensation-mediated N-WASP activation. We *de novo* engineered a trimeric SH3 protein by fusing SH3^FBP^^17^ with a well-defined trimeric coiled-coil motif ^53,54^ (Figure 5E). The recombinant trimer-SH3^FBP^^17^ was purified and incubated with N-WASP on Flat SLB. We first characterized N-WASP oligomerization by single-particle imaging via TIRFM. We found that the control monomeric SH3^FBP^^17^ doesn’t induce obvious clustering of N-WASP, while trimerCC-SH3^FBP^^17^ resulted in higher signal intensity in N-WASP clusters (Figure 5F,G). We then assessed whether the multivalent SH3 induces N-WASP clustering and activates actin assembly. Via TIRFM actin polymerization assay, we compared actin spontaneous polymerization using auto- inhibited N-WASP FL with Arp2/3 complex, with using monomeric or trimeric SH3 domain. A significant increase in F-actin production and the formation of star-shaped actin clusters were observed in the presence of trimer-SH3^FBP^^17^ but not in monomeric-SH3^FBP17^, indicating a valency-dependent activation machinery of N-WASP FL (Figure 5H,I; Supplementary Figure S6A-B, Movie S3). To summarize, our results highlight the indispensable role of multivalent interaction to cluster and activate N-WASP via SH3 domain in curvature sensor FBP17 (Figure 6).

**Figure 6.**
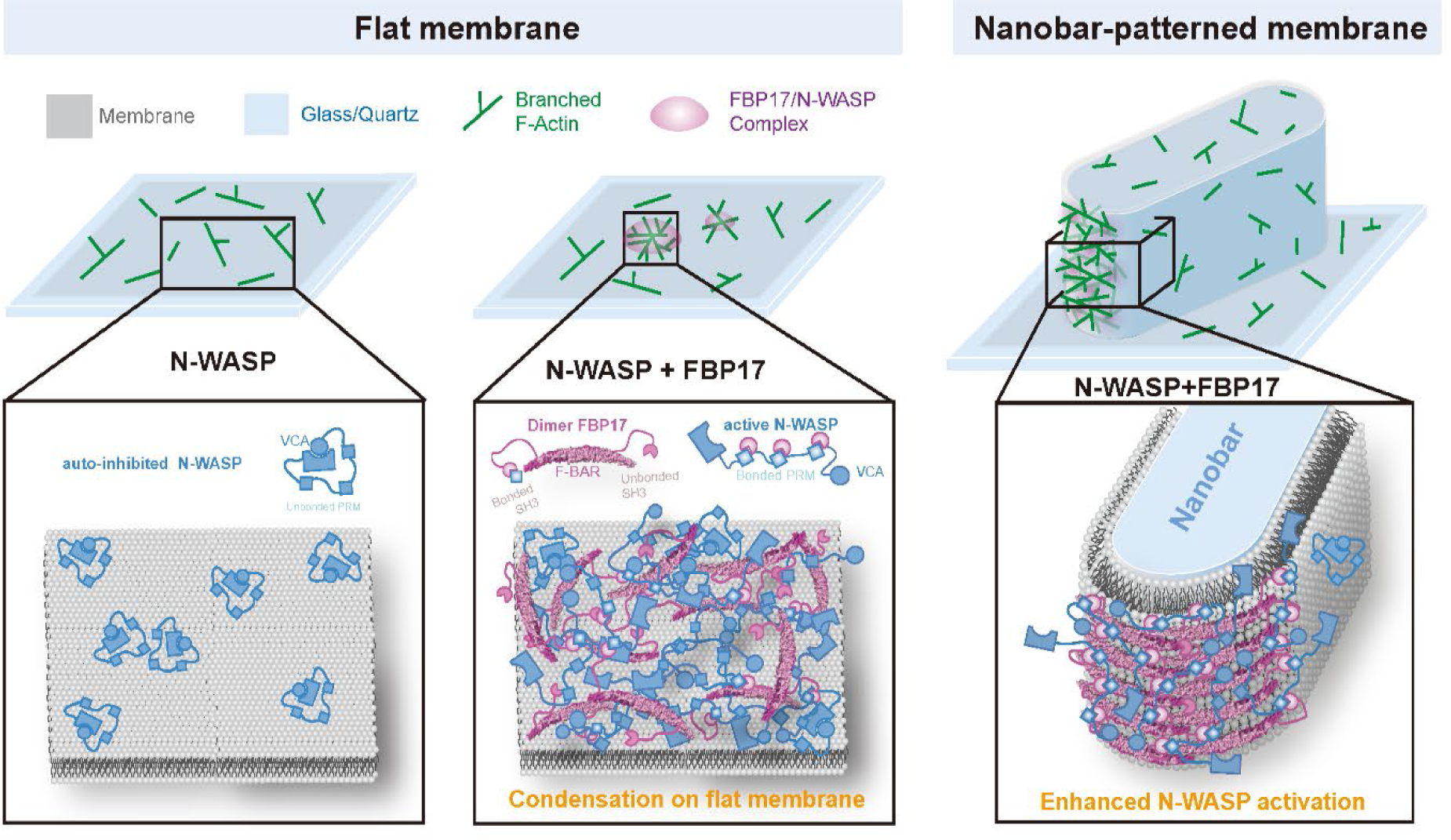
Proposed model of curvature-facilitated molecular condensation and F-actin polymerization around nanobar curved ends. On flat membrane surfaces, N-WASP molecules bind randomly with membrane (left panel); while the SH3 domains of FBP17 engage in multivalent interactions with N-WASP’s PRM, facilitating the exposure of VCA domains to promote branched F-actin polymerization (middle panel). Conversely, on nanobar-patterned membranes, FBP17’s F-BAR domain prefers the induced membrane curvature over flat surfaces, leading to the enrichment and local condensation of N-WASP. This process of molecular condensation significantly increases the occupancy of the polyP region, thereby enhances N-WASP activity in actin polymerization (right panel).

## Discussion

### Orchestrating Membrane Progression by Clustering Actin Nucleation Complex via Curvature Sensor BAR Protein

Our study applies minimal reconstitution systems and nanoscale modifications to lipid bilayers to map the regulation of actin polymerization by membrane curvature. We show how curvature influences the assembly of multicomponent systems like F-BAR and N-WASP, with the curvature radius fine-tuning actin nucleation. This underlines a complex relationship between membrane shape and the initiation of actin networks, where protein recruitment and activity are tuned by curvature. Our research reveals how membrane curvature regulates actin assembly spatiotemporally, influencing and being influenced by actin dynamics. Membrane curvature not only responds to forces but directs actin dynamics, which in turn sculpt the membrane topography, driving processes like endocytosis and producing cellular patterns such as oscillations ^7,31,55–58^. This interplay is evident in phenomena like PM wave propagation, where the dynamics of BAR domain-containing proteins align with curvature changes, suggesting a regulatory relationship where curvature guides actin polymerization to alter PM topography ^8,58^.

In particular, the F-BAR domain’s association with N-WASP is critical in membrane invagination ^31^, showing a precise response to curvature (Figure 1). F-BAR proteins, exemplified by TOCA-1, FBP17 and CIP4, display an affinity for mild positive curvatures with their extended BAR domain structure. There F-BAR proteins coordinate with other proteins in curvature-related processes such as clathrin-independent endocytosis and the formation of membrane tubules ^32,59^, suggesting a role for F-BAR-mediated multivalency. *In vitro* studies have shown the SH3 domain of these proteins can directly activate N-WASP-mediated actin polymerization on liposomes with a specific diameter ^21,26^. The other BAR proteins, like I-BAR (IRSp53), recruit actin regulatory proteins like VASP, mDia2 and WAVE2 through SH3 domain interaction ^27,60–67^. Our research now clarifies how the SH3 domain contributes to nanoscale N-WASP clustering upon precise curvature sensing by BAR, promoting localized Arp2/3 activation and robust actin nucleation.

### Spatiotemporal Coordination of Rapid Nano-scale Assembly and Activation of Actin Nucleation Complex on Plasma Membrane

Our research elucidates an additional mechanism of N-WASP activation at highly curved membrane region, that operates independently of the well-characterized Cdc42 pathway. The auto-inhibited state of N-WASP is typically relieved via a conformational release facilitated by Cdc42 and PI(4,5)P2, enabling interaction with the Arp2/3 complex and subsequent actin polymerization ^35,36^. However, alternative modes that enable N-WASP/Arp2/3 complex activation have also been observed, although the mechanism was unclear, particularly in multicomponent systems. For example, recombinant N-WASP displayed inherent actin polymerization activity, evident in both *in vitro* actin assays and cellular extracts, without relying on Rho-GTPase ^68^. Cellular extracts, however, seemed to lack a sturdy, spatially-regulated activation mechanism associated with Rho GTPase switch activation, typically contingent on membrane recruitment ^69,70^.

Here, we demonstrate that N-WASP can be activated through multivalent interactions with the F-BAR protein FBP17, mediated by PRM-SH3 binding, independent of Cdc42. This alternative pathway, which leads to the clustering of N-WASP, achieves localized activation around curvature without the direct involvement of Rho-GTPase. It underscores a multifaceted control of actin nucleation, further substantiated by phase separation studies that suggest a stoichiometry- and dwell-time-dependent enhancement of actin nucleation ^39,40,71,72^. Such condensation-driven activation events have also been observed in other systems. In plants, the activation of formin nucleators, single transmembrane domain proteins situated on the PM, were also found to rely on nano-scale condensation during immune responses and bypasses the need for a GTPase-regulated mechanism ^72,73^. This mechanistic diversity in regulating actin nucleation factor complex could have broad implications for the spatiotemporal organization of the cytoskeleton in cellular processes. First, membrane curvature acts as a catalyst, precisely orchestrating the spatial arrangement of proteins essential for actin polymerization. It shapes a unique three-dimensional platform, enhancing recruitment and concentration of BAR-domain proteins and N-WASP, resulting in functional clusters primed for actin assembly, which could happen in a Cdc42-independent manner, thought it is likely still coupled with Cdc42- activation machinery *in vivo*. Second, the topography feedback regulates component stoichiometry by the conformation and availability of N-WASP polyP and the FBP17 SH3 domain. These are governed by the assembly pattern of FBP17, dictated by BAR-domain-mediated sensing of topographical cues. When the system initially promotes interaction and assembly of the linker protein FBP17 in regions of curvature, it orients FBP17 array to present its SH3 domain while concealing other potential binding sites that could lead to non-specific associations. It ensures quick condensation of N-WASP in curved areas through SH3-PolyP interactions. As the optimal stoichiometry is achieved through a slim layer of the FBP17 array, the availability of the FBP17 SH3 domain decreases in proportion to the expanding condensation of N-WASP on the surface. When situated on flat membranes, where there are no curvature-defined array patterns for FBP17, the localized gatherings of the N-WASP and FBP17 complex progressed more slowly. Stoichiometry progressed more linearly as these biomolecules could expand in various directions, unrestricted by surface curvature. Such interplay between topographical cue and molecular assembly might also contribute to the observed dynamic behavior of molecules observed during membrane invagination and the oscillatory patterns that coincide with changes in topography ^57,74,75^.

### Limitations of the Study

The modulation of N-WASP activity by FBP17 clustering, whether by altering intramolecular interactions as suggested for Toca-1 ^26^ or by instigating direct conformational changes to expose the C-terminus, remains to be conclusively delineated by structural studies. Adding to the complexity is the challenge of understanding how nano- scale curvature determines N-WASP’s biochemical activity in living cells—a promising yet experimentally complex area for future research.

The modeling strategies used in this study do not consider the fine molecular details of FBP17 or N-WASP. Further, the Shannon entropy framework does not consider enthalpic interactions between N-WASP molecules. These are areas for future research that would be crucial to understand the role that molecular interactions play in amplifying actin signaling.

Cellular membrane curvature varies widely, spanning from tens to thousands of nanometers. Electron beam lithography (EBL) currently faces limitations in detailing features below 100 nm radius, especially with high aspect ratios, due to the proximity effect and etching precision. However, the use of nanostructured systems to create controlled curvature gradients now allows for the study of protein interactions specific to curvatures up to 100 nm radius. This capability to observe the dynamics between biomolecules and nanoscale curvature enriches our investigative potential. With improvements in EBL resolution and etching techniques, there’s potential to document a wider range of curvature scales, matching the complete spectrum found in cells. Integrating these advances with high-resolution imaging is set to significantly enhance our understanding of biomolecular assembly.

## Resource Availability

### Lead Contact

Further information and requests for resources and reagents should be directed to the Lead Contact, Yansong Miao.

### Materials Availability

All materials generated from this study are available upon request.

### Data and code availability

This study did not generate a new databset or code. Input files used for the simulations along with the analysis scripts will be available at https://github.com/RangamaniLabUCSD/FBP_NWASP_Nanopillar upon publication.

## Acknowledgments

We thank Dr. Min Wu (Yale University, USA) and Dr. Mingjie Zhang (SUSTech, China) for sharing plasmids for sub-cloning and protein production. This work was funded by the Singapore Ministry of Education (MOE) Academic Fund Tier 3 (MOE2019-T3-1-012 to Y.M. MOE-MOET32020-0001 to W.Z.), Tier 2 (MOE-T2EP30220-0009 and T2EP30222-0043

to W.Z.), and Tier 1 (RG95/19 and RG93/22 to W.Z.); National Research Foundation Singapore under its Open Fund - Individual Research Grant (MOH-000955 to Y.M.), National Research Foundation Singapore (NRF-NRFI08-2022-0012 to Y.M.), National Institutes of Health, NIGMS R01-GM132106 to P.R., and the Human Frontier Science Program Foundation (RGY0088/2021 to W.Z.), IDMxS to Y.M. and W.Z. and the Start-up Grant from Nanyang Technological University to W.Z.). The authors also acknowledge the support of EEE N2FC, SPMS CDPT, MSE FACTS for the fabrication and characterization of nanochips.

## Author Contributions

Conceptualization, K.Z., P.R., W.Z., and Y.M.; Methodology, K.Z., X.G., X.M., and A.C.;

Investigation, K.Z., X.G., and A.C.; Software, X.M. and A.C.; Formal Analysis, K.Z. and A.C.; Resources, X.G.; Writing – Original Draft, K.Z., X.G., A.C., P.R., W.Z., and Y.M.;

Writing – Review & Editing, K.Z., Y.M., A.C., P.R., X.G., X.M., and W.Z.; Funding

Acquisition, P.R., W.Z., and Y.M.; Supervision, P.R., W.Z., and Y.M.

## Declaration of Interests

The authors declare no competing interests.

## MATERIALS AND METHODS

Detailed methods are provided in the online version of this paper and include the following:

## METHOD DETAILS

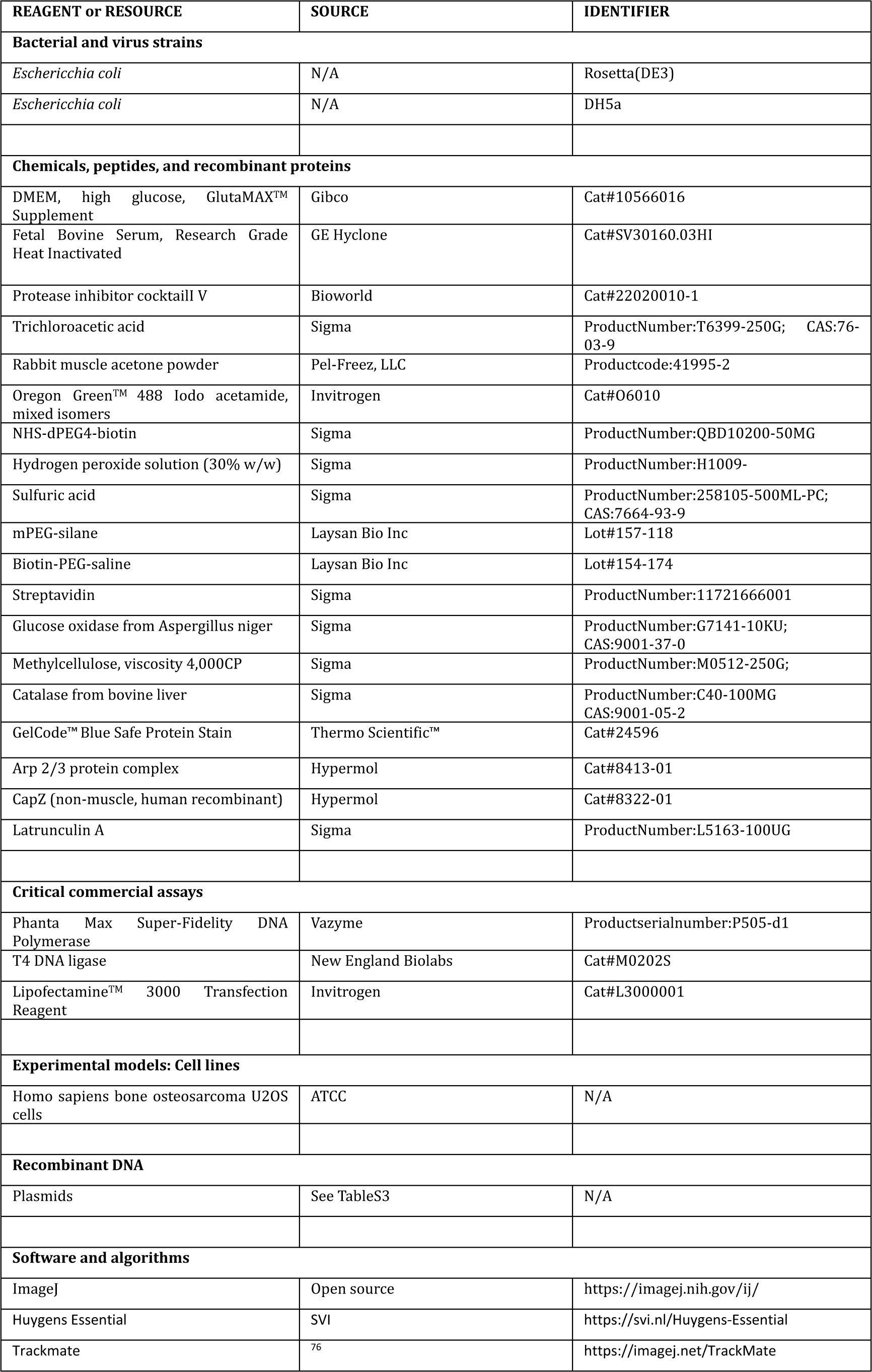
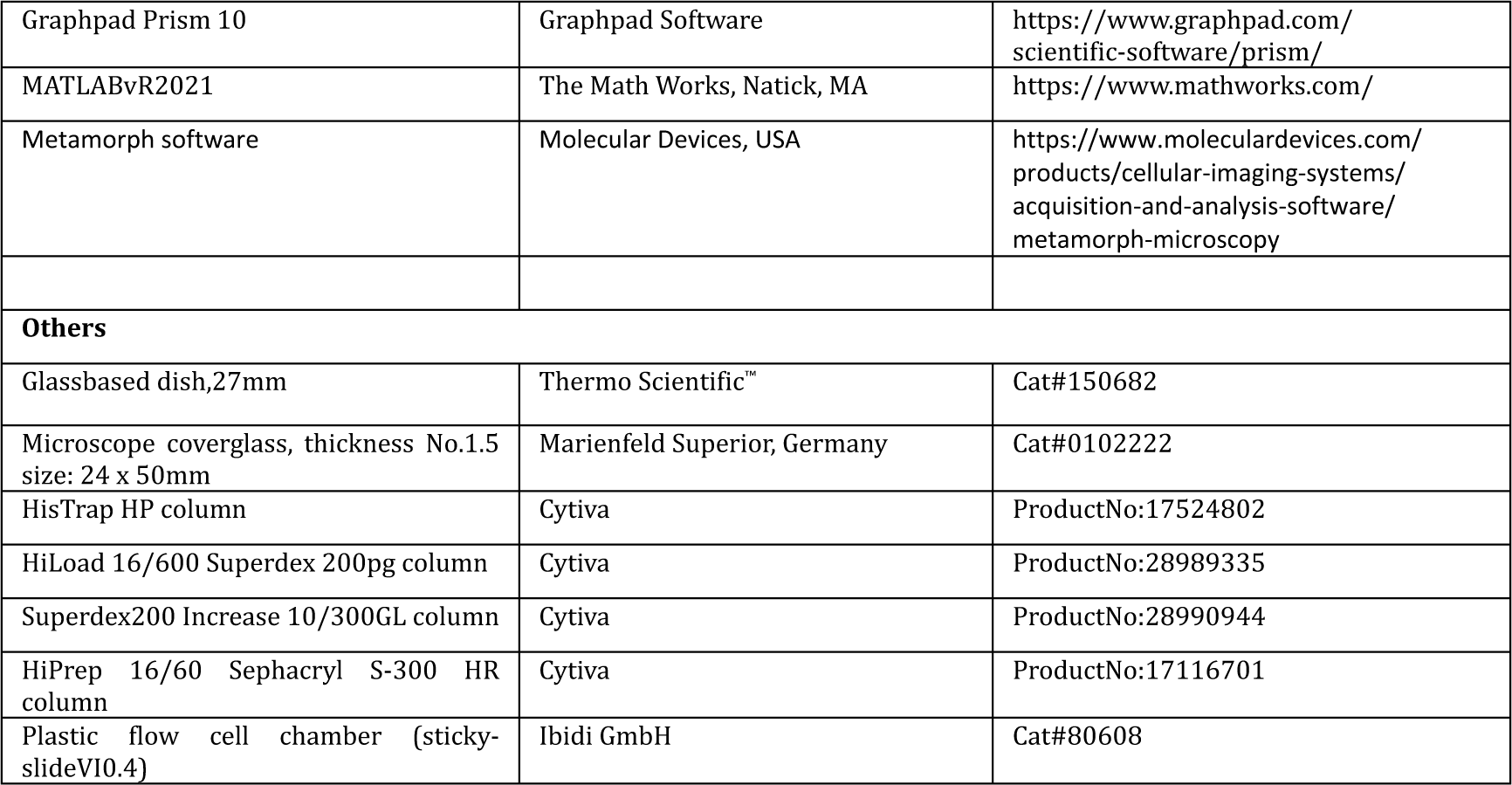

## **1.** Protein Purification and Fluorescent Labelling

### **1.1.** Protein reagents

The proteins FBP17 (aa 1-617) with an N-terminal His6 tag, N-WASP (aa 1-505), N-WASP L235P (aa 1-505), polyp VCA (aa 151-505), VCA (aa 392-505) with an N-terminal His6 tag, monomer-SH3, Trimer-SH3 with an N-terminal His6-Avi tag were produced and isolated from BL21(Rosetta) cells. The Arp2/3 complex and the CapZ protein was procured from Hypermol.

### **1.2.** FBP17, Monomer/Trimer-SH3 and N-WASP FL / L235P / 151-505 / VCA purification and labeling

BL21(DE3) Rosetta cells transformed with plasmids were selected using antibiotics. Subsequently, BL21 Rosetta cells were inoculated into 2 L of TB medium using a 50 mL overnight culture. The cells were then grown at 37°C until they reached an optical density (O.D.) of 1–2 at 600 nm absorbance, and protein expression was induced with isopropyl- thio-β-D-galactoside (IPTG, 1 mM) at 16°C for 16 hours. After induction, cells were harvested by centrifugation and resuspended in 45 mL of Binding Buffer (20 mM HEPES, pH 7.4, 500 mM NaCl, 20 mM Imidazole). Cell disruption was achieved using a homogenizer (LM20 Microfluidizer™) in Binding Buffer supplemented with 1 mM PMSF, 0.1% (v/v) Triton X100, and protease inhibitor cocktail tablets. The supernatant was obtained by centrifugation at 20,000 rpm for 1.5 hours at 4°C, filtered (0.22 μm), and then applied onto a 5 mL HisTrap HP column (G.E. Healthcare) connected to an FPLC AKTA system (G.E. Healthcare). The column was washed with Washing Buffer (20 mM HEPES, pH 7.4, 50 mM imidazole, and 500 mM NaCl), and proteins were eluted with Elution Buffer (20 mM HEPES, pH 7.4, 500 mM imidazole, and 500 mM NaCl). The proteins were further purified by size exclusion chromatography on a HiLoad 16/600 Superdex 200pg column (G.E. Healthcare) in Gelfiltration Buffer (20 mM HEPES pH 7.4, 500 mM NaCl, 10% Glycerol, and 1 mM DTT), and finally concentrated using 15 mL 50 kDa cut-off concentrators (Amicon Inc.) to a concentration of ∼5 mg/mL.

The primary amine group of FBP17, Monomer/Trimer-SH3, and N-WASP FL / L235P / CRIB-PolyP-VCA / VCA were labeled using an Alexa Fluor™647 and Alexa Fluor™488 labeling kit (Thermo Scientific). Briefly, the protein of interest was prepared at ∼2 mg/mL in a buffer containing 0.1 M sodium bicarbonate buffer. The Alexa dye was added to the mixture and incubated overnight at 4°C. Excess dye was removed by using a 5 mL HiTrap Desalting column (G.E. Healthcare) in Gelfiltration Buffer (20 mM HEPES pH 7.4, 500 mM NaCl, 10% Glycerol, and 1 mM DTT). Labeling efficiency was determined using a Nanodrop 2000 (Thermo Scientific).

### **1.3.** Rabbit skeletal muscle actin purification and labeling

To obtain monomeric ATP-bound rabbit muscle actin (RMA) for the SLB actin reconstitution assay and TIRF actin assembly assay, two grams of rabbit muscle acetone powder (Pel-Freez, LLC) were dissolved in 200 mL of cold G-buffer (2 mM Tris, pH 8.0, 0.2 mM ATP, 0.5 mM DTT, and 0.1 mM CaCl2) and stirred at 4°C overnight. Subsequently, the mixture was filtered with cheesecloth to remove muscle powder, and the actin- dissolved solution underwent centrifugation at 2600 x *g* and 4°C (Type 45 Ti rotor, Beckman Coulter) for 30 minutes to collect the supernatant. Actin in the supernatant was then polymerized with slow stirring for 1 hour at 4°C by adding KCl and MgCl2 solution to final concentrations of 50 mM and 2 mM, respectively. To remove tropomyosin and other actin-binding proteins, fine KCl powder was slowly added to reach a final concentration of 0.6 M, and the solution was stirred for another 30 minutes. The solution was subsequently centrifuged at 14,000 x *g* (Type 45 Ti rotor, Beckman Coulter) for 3 hours at 4°C to collect the filamentous actin pellet. The pellet was then rinsed with cold G-buffer and homogenized with a homogenizer in 7 mL of cold G-buffer, followed by a short sonication time.

The sample was then dialyzed in 2 L G-buffer at 4°C for 48 hours to induce depolymerization, with G-buffer changed every 12 hours. After buffer exchange, the sample underwent centrifugation at 167,000 x *g* (S.W. 55 Ti swinging-bucket rotor, Beckman Coulter) at 4°C for 2.5 hours, and 5 mL of supernatant was collected and loaded onto a Sephacryl S-300 HR column (G.E. Healthcare) pre-balanced with G-buffer. Peak fractions were collected and combined, and then 0.01% (final) sodium azide (Sigma) was added to inhibit fungal contamination, maintaining the sample at 4°C. Actin concentration was measured by assessing the OD290 with a Nanodrop 2000 (Thermo Scientific).

For actin labeling with Oregon GreenTM 488 iodoacetamide (Invitrogen), the same purification steps were followed as for RMA until the pelleted filamentous actin was homogenized and sonicated. After this, the sample was dialyzed in 1 L G-buffer at 4°C overnight. The next day, the sample was changed to 1 L G-buffer without DTT and dialyzed for 4 hours at 4°C, with buffer changed once. Oregon GreenTM 488 iodoacetamide was dissolved in dimethylformamide to a final concentration of 10 mM. Before labeling, actin concentration was measured by reading the OD290 with a Nanodrop 2000 (Thermo Scientific).

Actin was first diluted with an equal volume of cold 2X labeling buffer (50 mM imidazole, pH 7.5, 200 mM KCl, 0.6 mM ATP, and 4 mM MgCl2) and further diluted to a final concentration of 23 mM with cold 1X labeling buffer. Then, a 10-fold molar excess of Oregon Green™ 488 iodoacetamide was added dropwise while very gently vortexing. The mixture was covered with aluminum foil and rotated at 4°C overnight. The next morning, labeled filamentous actin was centrifuged at 167,000 x *g* (Type 50.2 rotor, Beckman Coulter) for 3 hours at 4°C. Pellets were collected, homogenized in 4 mL G-buffer, left on ice for 1 hour, and homogenized again. Actin was then dialyzed in 1 L G-buffer at 4°C for 48 hours to induce depolymerization (in darkness, with G-buffer changed every 12 hours). After buffer exchange, actin was centrifuged at 436,000 x *g* (TLA100 rotor, Beckman Coulter) at 4°C for 1 hour. The supernatant was collected and further purified by a Sephacryl S-300 HR column (G.E. Healthcare) pre-balanced with G-buffer. Peak fractions were collected and combined, then dialyzed in 500 mL G-buffer with 50% (v/v) glycerol at 4°C overnight to reduce volume. Small aliquots were frozen in liquid nitrogen and stored in a -80°C freezer.

## **2.** Preparation of Supported Lipid Bilayer on Nanobar Chips

### **2.1.** Fabrication of nanobar chips

The nanobar arrays utilized in this study were fabricated on square quartz coverslips using electron-beam lithography, as described by ^9^. Briefly, 15×15×0.2 mm square quartz coverslips were initially spin-coated with positive electron-beam resist polymethyl methacrylate (Allresist) to a height of approximately 300 nm. A conductive protective coating was subsequently applied using AR-PC 5090.02 (Allresist). Nanobar patterns were then delineated via Electron Beam Lithography (F.E.I. Helios NanoLab) and developed in the developer AR 600-56 (Allresist). A 300 nm-height chromium mask was generated through thermal evaporation (UNIVEX 250 Benchtop) and lifted off using acetone. The nanobars were then created via reactive ion etching with a mixture of CF4 and CHF3 (Oxford Plasmalab80). Following the application of a 10 nm Cr layer, the nanostructures underwent characterization through scanning electron microscopy to examine their curvature diameter and structural height. Before use, the nanobar chips underwent immersion in Chromium Etchant (Sigma-Aldrich) until the chromium masks were completely removed.

### **2.2.** Preparation of lipid vesicles

The lipid vesicles were composed of 1-palmitoyl-2-oleoyl-glycero-3-phosphocholine (POPC) mixed with 0.5 mol% of 18:1 Rhodamine-PE, 1 mol% of phosphatidylinositol 4,5- bisphosphate (PI(4,5)P2), and 10 mol% of 1-palmitoyl-2-oleoyl-sn-glycero-3-phospho-L-serine (POPS). Initially, the lipids were dissolved in chloroform and mixed at the specified molar ratios. The lipid mixture was then dried down in a brown glass vial using 99.9% nitrogen gas for 5 minutes, followed by vacuum drying for 1.5 hours to eliminate residual chloroform. Subsequently, the dried lipid film was resuspended in phosphate-buffered saline (PBS) at a concentration of 0.5 mg/mL and sonicated for 30 minutes. The resulting lipid mixture was transferred to a 1.5 mL tube and subjected to freeze-thaw cycles for 15 times (20 seconds in liquid nitrogen followed by 2 minutes in a 42°C water bath). The lipid mixture was further extruded through a polycarbonate membrane with a pore size of 100 nm using an extruder equipped with a holder/heating block (610000-1EA, Sigma-Aldrich). Finally, the lipid vesicle solution was stored at 4°C and utilized within 7 days.

### **2.3.** Formation of protein-bound supported lipid bilayers (SLBs) on nanobar chips

The nanobar chips underwent cleaning with piranha solution (composed of 7 parts concentrated sulfuric acid and 1 part 30% hydrogen peroxide solution) overnight, followed by rinsing with a continuous stream of deionized water to eliminate the acids. Subsequently, the nanobar chips were dried using 99.9% nitrogen gas and subjected to cleaning with air plasma in a plasma cleaner (Harrick Plasma) for 1 hour to eliminate any remaining impurities on the surfaces prior to lipid bilayer formation. Once cleaned, the chips were attached with a polydimethylsiloxane (PDMS) chamber, and lipid vesicles were introduced into the PDMS channel, where they were allowed to incubate for 15 minutes to facilitate the formation of the lipid bilayer. PBS was employed to rinse away any unbound vesicles within the chamber. Following this, protein solution of desired concentration was premixed with the indicated molar ratios in PBS and incubated for 2 min before introducing onto the lipid-bilayer-coated nanobar. Microscopy imaging was conducted after 30 minutes incubation at room temperature.

### **2.4.** Imaging of protein-nanobar interaction

The distribution of purified proteins on lipid-bilayer-coated nanobar arrays was investigated using a spinning disc confocal (SDC) system built around a Nikon Ti2 inverted microscope equipped with a Yokogawa CSU-W1 confocal spinning head and a 100X/1.4NA oil immersion objective. The Rhodamine-PE-containing lipid bilayer was excited at 561 nm and detected at 570-645 nm. Alexa Fluor 488-labeled N-WASP was excited at 488 nm and detected at 490-570 nm, while Alexa Fluor 647-labeled FBP17 was excited at 633 nm and detected at 645-700 nm.

## **3.** Reconstitution Assay on Flat SLBs

### **3.1.** Formation of protein-bound SLBs on flat surfaces

Flat SLBs were prepared in 96-well glass-bottomed plates (Matrical). Initially, wells were subjected to washing with 1 L of 5% (v/v) Hellmanex III (Hëlma Analytics) overnight while gently vortexed by a magnetic stirrer. Following this, the wells were thoroughly rinsed with deionized water (DI H2O) for 20 cycles, washed with 5M NaOH for 1 hour at 50°C for three repetitions, and again thoroughly rinsed with DI H2O. Subsequently, the wells were dried using 99.9% nitrogen gas and treated with air plasma in an HP plasma cleaner (Harrick Plasma) for 90 minutes to remove any remaining organic contaminations on the glass surfaces before SLB formation. SUVs were added into the wells and incubated for 15 minutes at 42°C to allow the SUVs to collapse and fuse onto the glass, thereby forming SLBs. The SLBs were then rinsed with PBS to wash away any unbound SUVs. To reconstitute the protein interaction on lipid membrane, protein solution of desired concentration was premixed with the indicated molar ratios in PBS and incubated for 2 min, and further added to the well. After 15 min of incubation, unbound proteins were removed by washing with PBS.

### **3.2.** Imaging of protein clustering on flat SLBs

Images were acquired at room temperature using a Nikon Ti2-E inverted microscope equipped with a 100x 1.45NA Plan-Apo objective lens and a TIRF module (iLasV2 Ring TIRF, GATACA Systems) and an ORCA-Fusion sCMOS camera (Hamamatsu Photonics). Image acquisition was controlled by MetaMorph software (Molecular Device).

## **4.** U2OS live-cell imaging

### **4.1.** Cell culture

Homosapiens bone osteosarcoma U2OS cells (ATCC) were maintained in DMEM with GlutaMAX (Gibco) supplemented medium with 10% FBS. and 1% PS. All cell lines were seeded in 100 mm plastic dishes with ∼1 × 10^6^ cells per dish. Cells were kept in an incubator with 5% CO2 and 100% relative humidity at 37 °C.

### **4.2.** Cell imaging with nanobar-substrates

To enable cell imaging on nanochip, the chip was attached to the 35 mm cell culture dish (TPP) with a hole punched in the center to expose nanobar pattern. Before cell plating, the dish substrate was sterilized by UV treatment for 20 min. The surface was coated with 0.2% Fibronectin (Sigma-Aldrich) for 30 min to promote cell attachment. U2OS cells were then cultured on the nanobar chip for one day before transfection. U2OS cells were then transiently transfected with the lifeact-mApple plasmids using Lipofectamine 3000 (Invitrogen, USA) following the manufacturer’s protocol and grown overnight on coated nanobar chip at 37 °C in a CO2 incubator for protein expression. 100 nM LatA or the same volume of DMSO was added to the cells and incubated for 1h at 37 °C and washed 3 times with PBS and changed back to DMEM. Cell imaging was then performed with a SDC built around a Nikon Ti2 inverted microscope containing a Yokogawa CSU-W1 confocal spinning head and a 100 X/1.4NA oil immersion objective. During imaging, the cells were maintained under 37 °C with 5% CO2 in an on-stage incubator.

## **5.** Actin TIRF assays and lipid-bound reconstitution assays

### 5.1. TIRF actin assembly assay *in vitro*

The *in vitro* real-time actin assembly assay was conducted on Biotin-PEG coated glass slices (Laysan Bio Inc) in 6-well chamber slices (Ibidi). The chamber was first blocked by 30 mL HBSA buffer (20 mM HEPES pH7.4, 1 mM EDTA, 50 mM KCl, 1%(m/v) BSA.) and incubated for 30 s. The glass surface was then conjugated with streptavidin by adding 30 mL HEKG10 buffer (20 mM HEPES pH7.4, 1 mM EDTA, 50 mM KCl,10%(v/v) glycerol, plus 0.1 mg/mL streptavidin) and incubating for 1 min. Afterward, free streptavidin was washed away using 1x TIRF buffer (10 mM imidazole pH 7.4, 50 mM KCl, 1 mM MgCl2, 1 mM EGTA, 50 mM DTT, 0.3 mM ATP, 20 nM CaCl2, 15 mM glucose, 100 mg/mL glucose oxidase, 15 mg/mL catalase, 0.25% methylcellulose). Next, 30 μL 3X protein-actin mix containing 1.5 μM G-actin (79% purified globular rabbit actin, 20% Oregon Green 488-actin, 1% Biotin-actin), 200 mM EGTA, 110 mM MgCl2, and desired protein solution was mixed with 30 μL 2X TIRF buffer and added into the chamber to a final volume of 90 μL to initiate the actin polymerization. Images were acquired as a stack at room temperature with 15- sec intervals for 15min using a Nikon Ti2-E inverted microscope equipped with a 100x 1.45NA Plan-Apo objective lens and a TIRF module (iLasV2 Ring TIRF, GATACA Systems) and an ORCA-Fusion sCMOS camera (Hamamatsu Photonics). Imaging lasers were provided 488 nm/150 mW (Vortran) and 639 nm/150mW (Vortran) combined in a laser launch (iLaunch, GATACA Systems). Focus was maintained by hardware autofocus (Perfect Focus System), and image acquisition was controlled by MetaMorph software (Molecular Device). To quantify the number of actin branches, 13x13 μm^2^ ROIs were chosen from each time point image and the actin filament branches in each ROI were manually counted.

### **5.2.** Reconstitution of actin polymerization on SLBs

SLBs was formed and coated with proteins as described above. After a pre-actin image was acquired, the SLBs were then rinsed by 60 μL 1X TIRF buffer for 3 times. Subsequently, 30 μL 4X actin mix containing G-actin (10% Oregon Green™ 488 labeled), Arp2/3 complex, 0.25 mM ATP, 0.1 mM MgCl2, 0.1 mM EGTA, and desired protein solution were added and mixed with 30 μL 1X TIRF buffer into the well with gentle pipetting. For all experiments, TIRF images were acquired at 15 s intervals for 15 min. Oregon Green 488- actin images were acquired using 100 ms exposure and the 488-laser power set to 15%. AF647-FBP17 images were acquired using 200 ms exposure and the 647-laser power set to 50%. No detectable crosstalk was observed with these settings.

## **6.** Modeling Curvature Sorting of FBP17 and N-WASP along a Nanobar

### **6.1.** Reaction-diffusion framework

To study the curvature-sorting of FBP17 and N-WASP proteins, we used the VCell Math Model setting (www.vcell.org) ^47,48^ to model and simulate the system as a set of coupled bulk-surface partial differential equations (PDEs). We approximate the system in two dimensions and represent the cross-section of the nanobar as a rounded rectangle with varying diameters (between 100 and 1000 nm, as shown in Table S1). Additionally, we approximate the flat sides of the nanobars as elliptical arcs. This is essential to prevent discontinuities in the curvature profile along the arc length. As the nanobar cross-sectional area scales with bar width, we scaled the total reaction area to ensure moles of FBP17 and N-WASP remain constant across all bar widths considered.

Further, we do not represent the membrane explicitly. Instead, based on experimental observations (Figure 1), we assume that the membrane curvature at any point matches the local curvature of the nanobar. FBP17 and N-WASP molecules can diffuse freely in the volume around the nanobar. We consider the following reactions to model the curvature sorting of FBP17.

### **6.2.** Reactions on the membrane

#### 6.2.1. Hydrophobic patches on the membrane

Equilibrium thermal fluctuations of the membrane lead to short-lived exposure of hydrophobic tails of the lipid bilayer ^77^. Curvature-sensitive proteins such as FBP17 bind through enthalpic interactions with such hydrophobic tails of the lipids in the outer leaflet of the membrane, leading to helix insertion ^43^. To model this, we consider the spontaneous appearance and disappearance of hydrophobic patches (H) on the membrane.

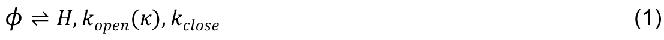

In addition, we assume that the rate of formation of defects depends on the local curvature of the membrane ^1^ using a curvature-dependent rate that is inspired by Bell’s law ^78^.

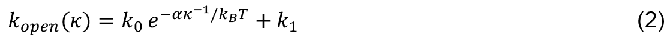

In the above equation, the rate of exposure of hydrophobic patches (𝑘_𝑜𝑝𝑒𝑛_) depends on the zero force opening rate (𝑘_0_), the zero-curvature binding rate (𝑘_1_) a characteristic force parameter (𝛼), the local curvature of the nanobar (𝜅), Boltzmann constant (𝑘_𝐵_) and temperature (𝑇). Additionally, the term (𝐹/𝜅) represents the energetic cost associated with the hydrophobic patch. The hydrophobic patches formed do not diffuse along the nanobar.

#### 6.2.2. FBP17 binding to hydrophobic patches

Diffusing FBP17 molecules bind to hydrophobic patches to stay bound to the nanobar.

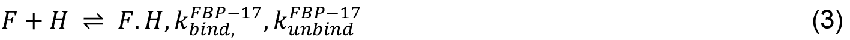

F.H molecules do not diffuse along the nanobar. To determine the kinetic parameters, we simulate FBP17 binding at various concentrations and compute the profiles of FBP17 nanobar-end density (Supplementary Figure S5 E). We see that at any given 𝑘_𝑏𝑖𝑖𝑛𝑑,𝐼_ value, increasing 𝑘_𝑢𝑛𝑏𝑖𝑖𝑛𝑑,𝐼_ beyond a certain value leads to poor sorting of FBP17 along the nanobar ends (For example, profiles corresponding to 𝑘^𝐹𝐵𝑃−17^ = 10.0/𝑠. We also see that at certain parameter values the transition from low nanobar-end density to high nanobar-

end density is drastic (For example, 𝑘^𝐹𝐵𝑃−17^ = 100 𝜇𝑚^2^/(𝑚𝑜𝑙𝑒𝑐𝑢𝑙𝑒. 𝑠), 𝑘^𝐹𝐵𝑃−17^ < 10.0/𝑠).

𝑏𝑖𝑖𝑛𝑑

Ignoring such parameter pairs, we chose (𝑘_𝑏𝑖𝑖𝑛𝑑_, 𝑘_𝑢𝑛𝑏𝑖𝑖𝑛𝑑_)=(10.0,1.0).

#### 6.2.3. N-WASP binding to membrane-bound FBP17

𝑢𝑛𝑏𝑖𝑖𝑛𝑑

N-WASP molecules (N) that diffuse around the nanobar are tethered to the membrane through membrane-bound FBP17 molecules.

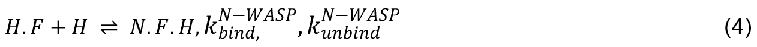

N.F.H molecules do not diffuse along the nanobar.

Please refer to Table S1 for a detailed description of the parameters used in the model.

### **6.3.** Geometric parameters

#### 6.3.1. Representation of the nanobar

The ideal representation of the nanobar shape would be a rounded rectangle. In our simulations, the nanobar-ends are represented by semi-circular ends whose diameter matches the bar width. Representing the flat sides of the nanobar as straight lines would lead to numerical inconsistencies due to abrupt changes in curvature at transition points as curvature goes from 1/R, R=bar width/2 to 0. Therefore, we modeled the flat sides as arcs of ellipses. The minor axis span, 2b was set to be 1.01% of bar width for bar width of 100 nm and as 1.05%. These choices were made empirically to ensure a less drastic curvature change. Assuming one of the nanobar semi-circles is centered at the origin, the ellipse major axis span 2a was determined by solving for the equation of the ellipse passing through (0, R).

#### 6.3.2. Reaction area

The nanobar area changes with bar width. Therefore, to ensure the number of moles of FBP17 and N-WASP available remains the same across all bar widths, we calculated the reaction area as follows. The length of the reaction area was kept constant at 2.5 𝜇𝑚 across all bar widths. Choosing a bar width of 1000 nm as a reference and the corresponding reaction area to be 3 𝜇𝑚^2^, we calculated the reaction area width (W_𝑟𝑥𝑛_). This gave a diffusible area of 2.2146 𝜇𝑚^2^. For all other bar widths, the reaction area width was calculated to ensure that the diffusible area is held constant at 2.2146 𝜇𝑚^2^.

### **6.4.** Modeling actin nucleation under varying N-WASP localization

We hypothesize that the localization of N-WASP leads to spatially-localized activation of Arp2/3 resulting in enhanced nucleation of Arp2/3. Thus, we test whether spatial localization of N-WASP is adequate to enhance nucleation activity using an agent-based modeling approach. In this model, we do not represent the membrane explicitly. Instead, we specify a spatial distribution of N-WASP concentration as the initial condition and study the role it plays in actin nucleation. Towards this, we first develop a model based on Shannon Entropy to systematically generate spatial copy number maps at progressively increasing N-WASP localization. Then, we simulate the resulting actin nucleation using Mechanochemical Dynamics of Active Matter v5.1.0 (MEDYAN v5.1.0).

#### 6.4.1. Generation of N-WASP spatial copy number maps

For this purpose, we consider a 16x16x1 compartment space, each of size 500x500x500 nm^3^ as the reaction volume. N-WASP copy number in each compartment can be specified at various levels of localization. Such copy number maps can be readily converted to probability density functions. Thus, to quantify spatial localization, we resort to the Shannon Entropy function.

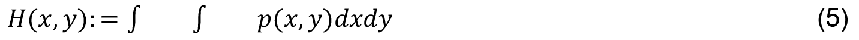

In the above equation, 𝑝(𝑥, 𝑦) represents the probability density function. Uniform probability density functions (represent random distribution of N-WASP without any localization) correspond to the maximum values of 𝐻(𝑥, 𝑦) = 𝐻_𝑚𝑎𝑥_. To compute 𝐻_𝑚𝑎𝑥_, we generate uniform distribution as the initial condition and the corresponding Shannon Entropy 𝐻_𝑚𝑎𝑥_. As the localization of N-WASP (p(x,y)) increases, the corresponding Shanon Entropy decreases (𝐻(𝑥, 𝑦) ≤ 𝐻_𝑚𝑎𝑥_). To generate such probability densities, we use the Sequential Least Squares Programming (SLSQP) algorithm specified in Python3.0 scipy.optimize.minimize to minimize the following objective function.

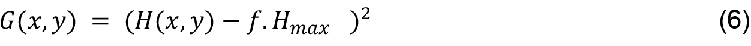

In the above equation, 𝑓𝑓 ∈ (0,1,] represents a factor of localization. When 𝑓𝑓 = 1.0, the minimum 𝐺_𝑚𝑖𝑖𝑛_ corresponds to an uniform distribution. At lower values, the corresponding minimum yields probability density distributions with increased localization. In this study, we consider 𝑓𝑓 values of 1.0, 0.99, 0.95, 0.8, and 0.5, respectively.

The resulting probability density functions are sampled to compute copy number maps corresponding to Arp2/3 activator molecule, N-WASP. To ensure that the resulting copy number maps give us the same Shannon Entropy values as the probability density functions, we progressively increase the total concentration and compute the relative error in Shannon entropy between the optimal density function and the sampled copy number map. We find low relative error values (<1%) at concentrations above 1𝜇𝑀. We consider the total N-WASP concentration to be 1𝜇𝑀 in this study. Please note that while these values do not correspond to experimental values of concentrations studied, the copy number, along with the rate of Arp2/3 activation, gives us a semi-quantitative simulation framework to study the role of localization.

#### 6.4.2. Mechanochemical Dynamics of Active Networks (MEDYAN)

To study actin nucleation resulting from the copy number maps, we use MEDYAN, a C++ stochastic mechanochemical simulator for actin networks. MEDYAN represents filaments explicitly as fibers and can also simulate the dendritic nucleation of actin filaments ^49^. We describe the chemical, mechanical, and mechano-chemical frameworks considered in this

study.

##### 6.4.2.1. Reaction-Diffusion Model

In MEDYAN, the reaction volume is divided into compartments of size 500 × 500 × 500 𝑛𝑚^3^. The copy number of chemical species is specified within each compartment. Stochastic diffusion is modeled as random hopping between neighboring compartments. In MEDYAN, reaction propensities are computed by assuming uniform mixing within each compartment. Thus, each compartment has reactions whose propensities are calculated based on the local concentration of reactive species in that compartment. Additionally, actin filaments are explicitly represented as fibers. We assume that every 27 nm of actin filament consists of one actin-binding site that is available for Arp2/3-driven nucleation. Thus, we consider the reactions shown in Supplementary Figure Y within each compartment.

While 𝐺 − 𝑎𝑐𝑡𝑎𝑎𝑛 and 𝐴𝑟𝑝2/3_𝑖𝑖𝑛𝑎𝑐𝑡𝑖𝑖𝑣𝑒_diffuse freely throughout the reaction volume, N-WASP molecules do not diffuse in our model. This assumption helps us capture the membrane-bound nature of N-WASP. In addition, as Arp2/3 activation is coupled with N-WASP binding, we assume that the activated Arp2/3 molecules do not diffuse freely.

##### 6.4.2.2. Next reaction method

The chemical evolution of the reaction network explained above is evolved in MEDYAN using a variant of the Gillespie method ^79^ called the *next reaction method* ^80^. Briefly, for every reaction 𝑅_𝜇_, 𝜇 ∈ [0, 𝑁) in the reaction network, we can compute reaction propensity

𝑎_𝜇_= 𝑐_𝜇_𝛾𝛾_𝜇_. Here, 𝑐_𝜇_represents the mesoscopic rate constant given by, 𝑐_𝜇_= 𝑘_𝜇_(𝑉_𝑟_/𝑁_𝐴_)^𝑛−1^, where 𝑘_𝜇_ is the rate constant of reaction of order 𝑛 happening in compartment of volume

𝑉_𝑟_and 𝑁_𝐴_represents the Avagardo constant. 𝛾𝛾_𝜇_represents the degeneracy of the reaction. For example, for the reaction 𝐴 + 𝐵 → 𝐶, 𝛾𝛾_𝜇_= 𝑁_𝐴_ × 𝑁_𝐵_ where 𝑁_𝐴_and 𝑁_𝐵_represent copy number of species A and B respectively within a given compartment. Once the propensities are computed, the timestep (𝑟_𝜇_) corresponding to each reaction can be computed randomly from 𝑟_𝜇_ = (1/𝑎_𝜇_)𝑙𝑛(1/𝑟_𝜇_), where 𝑟_𝜇_ represents a random number between 0 and

1. The reaction with the lowest 𝑟_𝜇_ is chosen and executed. Then, the subset of reactions 𝜇 that share reaction species with the executed reaction are chosen and the corresponding propensities and time steps are updated. Next reaction method speeds up this process by storing a dependency graph corresponding to the reaction network and also using an indexed priority queue data structure to efficiently sort the 𝑟_𝜇_ values. In MEDYAN, each of the chemical events are also coupled with a geometric change in the filament shape. For example, considering the monomer length to be 2.7 nm, (de) polymerization reactions lead to filament extension (reduction) by 2.7 nm. Additionally, dendritic nucleation reaction results in the formation of an offspring filament at 70° angle with respect to the parent filament. Thus, in addition to stochastic sampling, it is also essential to ensure MEDYAN generates mechanically equilibrated network architectures. To achieve this, MEDYAN uses a mechanical energy representation and an energy minimization protocol.

##### 6.4.2.3. Mechanical model

To ensure the physical realism of generated networks, in this study, we perform conjugate gradient energy minimization of actin networks after every 5 ms of reaction-diffusion perturbations.

#### 6.4.3. Representation of actin filament

Actin filaments are represented as a series of cylindrical fiber segments, each of length

𝐿_𝑐𝑦𝑙_ = 27 𝑛𝑚. As actin filaments resist extensile stresses, the segments are modeled as elastic springs with high spring stiffness (𝑘_𝑠𝑡𝑟_ = 100𝑝𝑁/𝑛𝑚). The stretching energy of a segment (i) is given by,

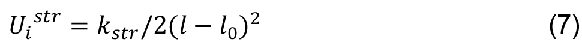

Here, the cylinder stretching constant is given by 𝑘_𝑠𝑡𝑟_, with current length 𝑙 and resting length 𝑙_0_. In this study, we choose a resting length 𝑙_0_=27 nm corresponding to 10 monomers.

Additionally, the filaments can bend along the hinge points according to the bending constant (𝑘_𝑏𝑒𝑛𝑑_) obtained from the experimentally measured flexural rigidity (𝐸𝐼 = 𝐿_𝑝_𝑘_𝐵_𝑇) of actin filaments. The bending constant is given by 𝑙_𝑐𝑦𝑙_𝑘_𝑏𝑒𝑛𝑑_ = 𝐿_𝑝_𝑘_𝐵_𝑇, where 𝐿_𝑝_,𝑘_𝐵_, and T represent persistence length of actin, Boltzmann constant, and temperature, respectively. In this study, we assume T=298K, and 𝑘_𝐵_𝑇=4.11 pN.nm. The corresponding bending energy of a hinge point between cylinders i and i+1 is given by,

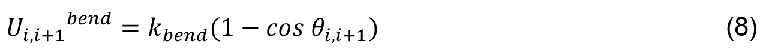

In addition, we explicitly model the steric repulsion to prevent spatial overlap of actin filament segments. Consider cylinders i and j. The excluded volume potential between the two is given by,

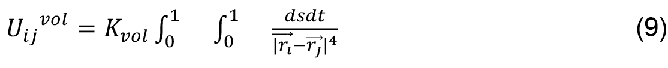

In the above equation, the excluded volume constant is given by 𝐾_𝑣𝑜𝑙_, and the distance between points on cylinder i and cylinder j is parameterized by representing the points on cylinder i with s and j with t. Please refer to Floyd et al. ^81^ for a detailed discussion on the excluded volume potential used here.

Finally, actin filament segments that are confined within the reaction volume through a repulsion potential with the planar boundary given as,

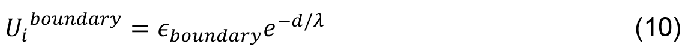

Here, the boundary repulsion depends on the distance from the boundary (d), boundary repulsion energy 𝜖𝜖_𝑏𝑜𝑢𝑛𝑑𝑎𝑟𝑦_and the screening length 𝜆 . Please refer to Supplementary Table S2 for a detailed description of the parameters used in this model.

##### 6.4.3.1. Dendritic potentials

The following potentials are considered between the parent and offspring filaments. To ensure that the parent and offspring filaments do not overlap and to ensure that the minus-end of the offspring filament remains at a definite position with respect to the parent filament, we consider a combination of a stretching and bending potentials. The length of bond connecting the binding site on parent and minus-end of offspring filament is given by, 𝐿_𝑏𝑜𝑛𝑑_ with a resting length 𝐿_0_ and is assumed to have a stretching constant 𝑘_𝑏𝑟𝑎𝑛𝑐ℎ,𝑠𝑡𝑟_.

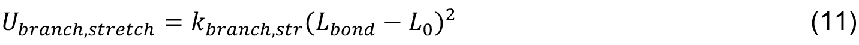

The angle (𝜃𝜃_𝑝𝑎𝑟𝑒𝑛𝑡,𝑏𝑜𝑛𝑑_) between the parent filament and bond is allowed to fluctuate around 𝜋/2 depending on the bending energy parameter 𝑘_𝑏𝑟𝑎𝑛𝑐ℎ,𝑏𝑒𝑛𝑑,𝐼_.

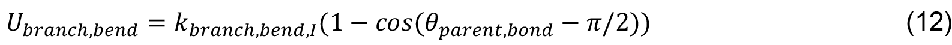

To ensure the angle between parent and offspring (𝜃𝜃_𝑝𝑎𝑟𝑒𝑛𝑡,𝑜𝑜𝑜𝑜𝑜𝑠𝑝𝑟𝑖𝑖𝑛𝑜𝑜_) is ∼70°, we consider a bending potential.

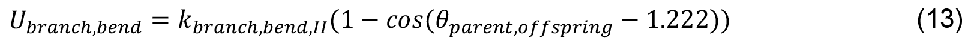

To penalize any twist along the bond connecting F-actin binding site and offspring minus-end, we consider a dihedral potential.

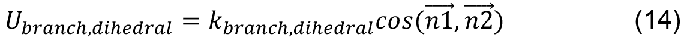

𝑘_𝑏𝑟𝑎𝑛𝑐ℎ,𝑑𝑖𝑖ℎ𝑒𝑑𝑟𝑎𝑙_ represents the dihedral energy parameter that penalizes the angle between plane normals (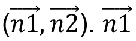 is the normal of plane encompassing by parent filament vector and the bond vector. 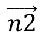 is the normal of plane encompassing the bond vector and the offspring filament vector.

6.4.3.2. Mechano-chemical models

The polymerization rate of filament ends that are close to the boundary is scaled according to the Brownian ratchet model ^82^.

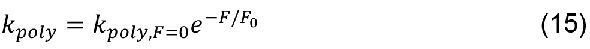

Hare, the polymerization rate under zero-force condition 𝑘_𝑝𝑜𝑙𝑦,𝐹=0_ of a filament end is modified to 𝑘_𝑝𝑜𝑙𝑦_ when it experiences force 𝐹 and a corresponding characteristic force 𝐹_0_=1.5pN.

## QUANTIFICATION AND STATISTICAL ANALYSIS

### Single particle intensity analysis

To quantify the intensity of AF488-N-WASP and AF647-FBP17 particles, the images were analyzed using Trackmate^76^ within ImageJ. Initially, a preliminary particle analysis was conducted manually, setting the estimated particle diameter to 0.65 μm (10 pixel) for particle detection. Additionally, quality control was performed to filter out false selections originating from the background, ensuring that only visible punctate protein signals were selected. The selected particles then underwent signal intensity analysis in Trackmate. Specifically, 200 out of more than 1000 particles from at least ten images were quantified for each protein combination to ensure statistical robustness.

### Curvature enrichment analysis of FBP17, N-WASP and actin signal on SLB

The processing and analysis of nanoarray images were conducted using custom-written MATLAB code adapted from a previous study ^42^. In brief, individual nanobar positions in the lipid channel image were located using a square mask (71×71 pixels) centered at the nanobar. The same mask was then applied to generate individual nanobar images of the protein channel. Background correction of each individual nanobar image was performed by subtracting a local background image generated using the meshgrid function in MATLAB based on four ROIs at the corners of the image. Background-corrected individual nanobar images with the same nanobar dimensions were averaged across different arrays and experimental repeats to generate averaged images. To quantify the signal at the nanobar-end and nanobar-center, each nanobar was segmented into three ROIs (two nanobar-ends and a nanobar-center). The size of the nanobar-end ROI was adjusted according to the dimension of the nanobar to minimize covering the nearby non-curved center of the nanobar. Intensity values of pixels within each ROI were then integrated.

For comparison between lipid and protein channels, the intensities of background-corrected individual nanobar images were normalized to the percentages of 600 nm nanobar-center intensity acquired in the same image. Protein density at each nanobar was measured by the ratio of normalized protein intensity to normalized lipid intensity. Additionally, montage of protein distribution on nanobars were obtained from averaged images using Fiji ImageJ. Statistical analysis was carried out using PRISM 10 (GraphPad).

### Curvature enrichment analysis of actin signal in U2OS cells

Background-corrected individual nanobar images with the same nanobar dimensions were averaged across different arrays and experimental repeats to generate averaged images as mentioned above. To quantify the signal at nanobar-end and nanobar-center, each nanobar was manually segmented into four ROIs (two nanobar-ends and a nanobar-center) as described in Figure 3C. Mean intensity values of pixels within each ROI were measured. The mean intensity at each nanobar-end was normalized to the averaged mean intensity of two centers for each nanobar size. Additionally, montage of lifeact-mApple signal on nanobars were obtained from averaged images using Fiji ImageJ. Statistical analysis was carried out using PRISM 10 (GraphPad).

### Statistics and reproducibility

All experiments are carried out with at least 2 replicates or otherwise indicated. All statistical analyses were done by GraphPad Prism (GraphPad). P values were determined by unpaired t-test assuming equal variances (*p< 0.05, **p< 0.01, -***p< 0.001, ****p< 0.0001, and ns = no significance).

## Supplemental Video

**Movie S1** Timelapse actin assembly on nanobar-patterned SLB

**Movie S2** TIRF actin assembly in the presence of N-WASP,FBP17 and CapZ

**Movie S3** TIRF actin assembly in the presence of SH3 monomer or SH3 trimer

---

**Supplementary Figure 1.**
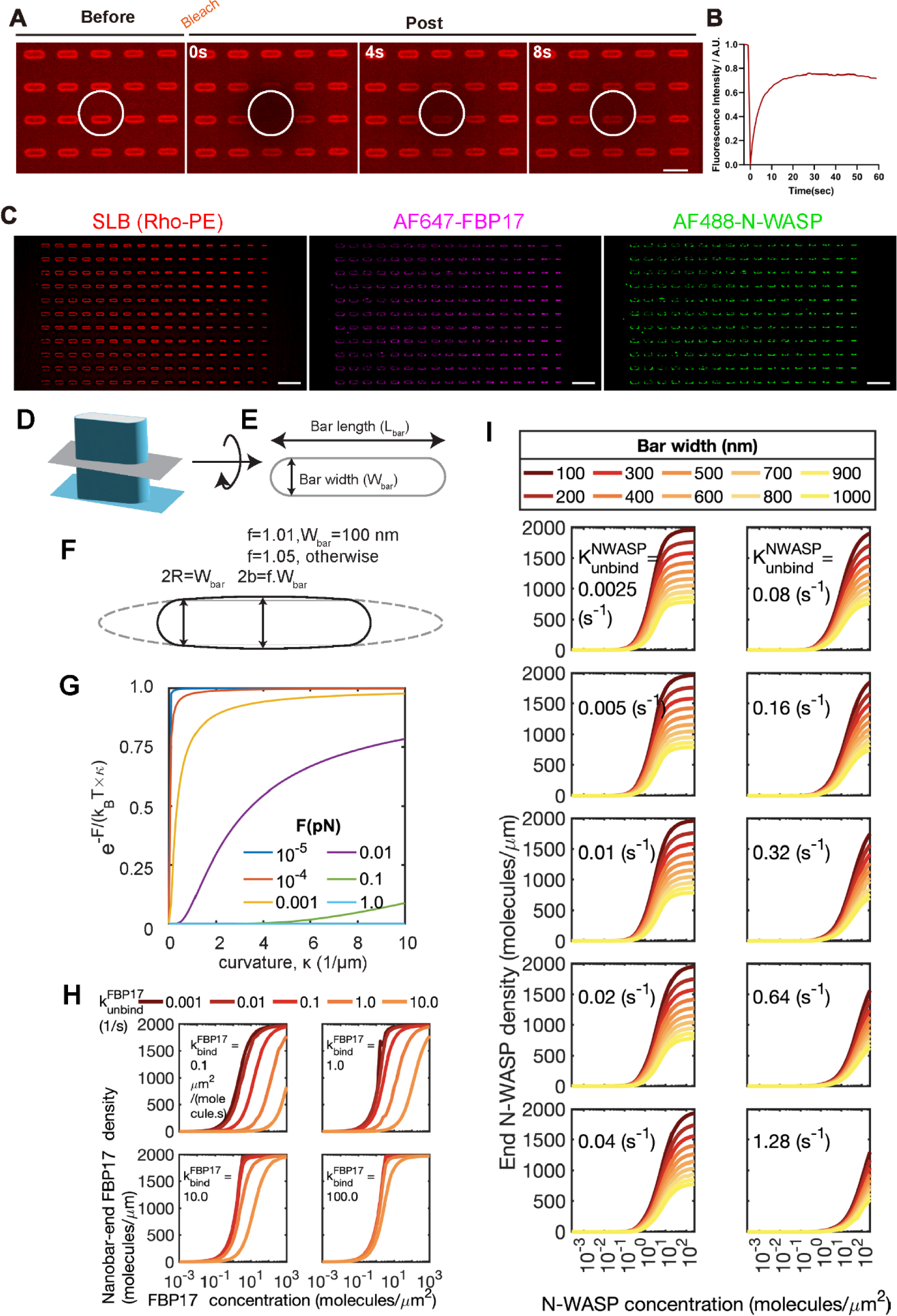
Distribution of N-WASP and FBP17 on dynamic patterned SLB. **A.** FRAP confocal images of dynamic SLB (10% POPS: 88.5% POPC: 1% PI(4,5)P2:0.5% Rhodamine-PE) on nanobar array. Scale bar, 5 μm. **B.** Normalized fluorescent signal recovery plot of SLB in **A**. **C.** Pre-averaged confocal images of SLB, AF647-FBP17 and AF488-N-WASP signal on curved SLB, 200-1000 nm bar width with 50 nm interval. **D**. Cartoon representation of a 3D nanobar, **E**. the cross-section (gray plane) of which is considered in our model. **F**. To prevent numerical instabilities, the nanobar (solid black line) is approximated as the union of an ellipsoid whose minor axis span (2b) scales depending on the bar width flanked by circular ends whose diameter matches the bar width. **G.** The force parameter in Eq (2) was chosen based on profiles of the force-sensitive term (Y- axis) as a function of local curvature (𝜅). **H.** Role of FBP17 binding and unbinding rates corresponding to 100nm bar width. The nanobar-end density of FBP17 molecules as a function of FBP17 concentration is plotted at various binding (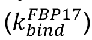) and unbinding rates (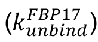). **I.** Role of N-WASP unbinding rate. Simulated profiles of nanobar-end density of N-WASP as a function of N-WASP concentration ([FBP17]:[N-WASP] = 4:1). 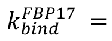 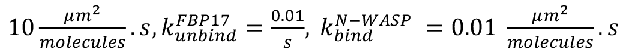 Each subplot is generated at a particular 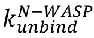 value and profiles are colored by the bar width.

**Supplementary Figure 2.**
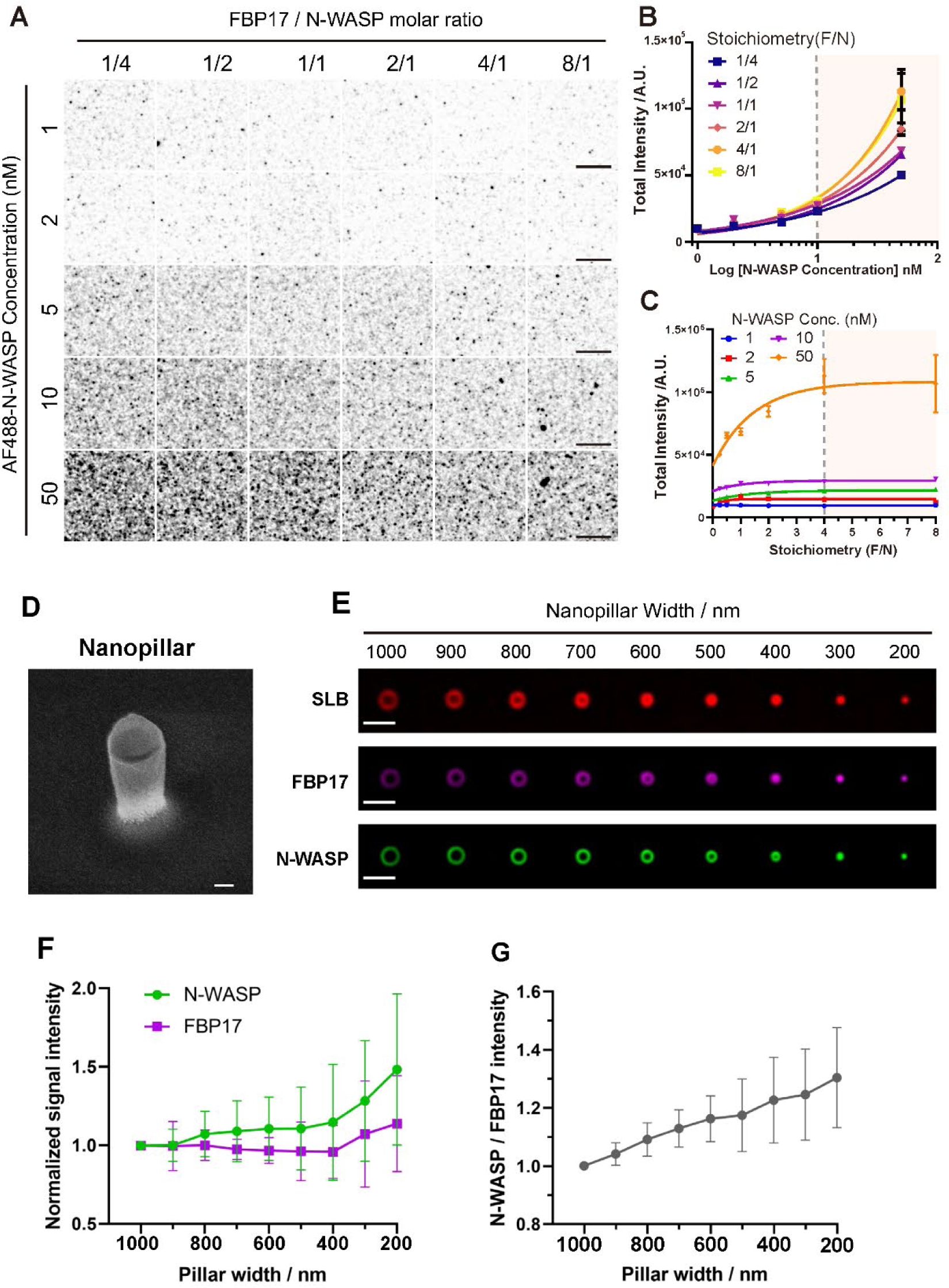
N-WASP forms condensate with FBP17 in a concentration- and stoichiometry-dependent manner. **A.** TIRFM single particle images of AF488-N-WASP at 1, 2, 5, 10, 50 nM on SLB with various stoichiometry of FBP17. Scale bar, 2 μm. **B.** Plot of single particle intensity of the N-WASP as a function of concentration on flat SLB in (**A**). Lines are binding curves fitted with the Hill equation. N=1000 from 3 repeated experiments, mean and SEM showed **C.** Total intensity plot of N-WASP single particles on SLB in (**A**). N=1000 from 3 biological repeats, mean and SEM are shown **D.** Scanning electron microscopy image of a single NanoPillar. Tilted 30°. Scale bar, 100 nm. **E.** Averaged confocal images of lipid bilayer (10% POPS: 88.5% POPC: 1% PI(4,5)P2:0.5% Rhodamine-PE) on top of nanopillars of 200-1000 nm width (upper panel), 200 nM AF647-FBP17 (middle panel), and 50 nM AF488-N- WASP (bottom panel) on the bilayer. Scale bar, 2 μm. **F.** Normalized nanopillar density of FBP17 and N-WASP based on their corresponding lipid bilayer intensity. Each point represents mean ± SEM from over 60 nanopillars. **G.** Normalized N-WASP end density divided by FBP17 end density at nanopillars of 200-1000 nm. Mean ± SEM showed.

**Supplementary Figure 3.**
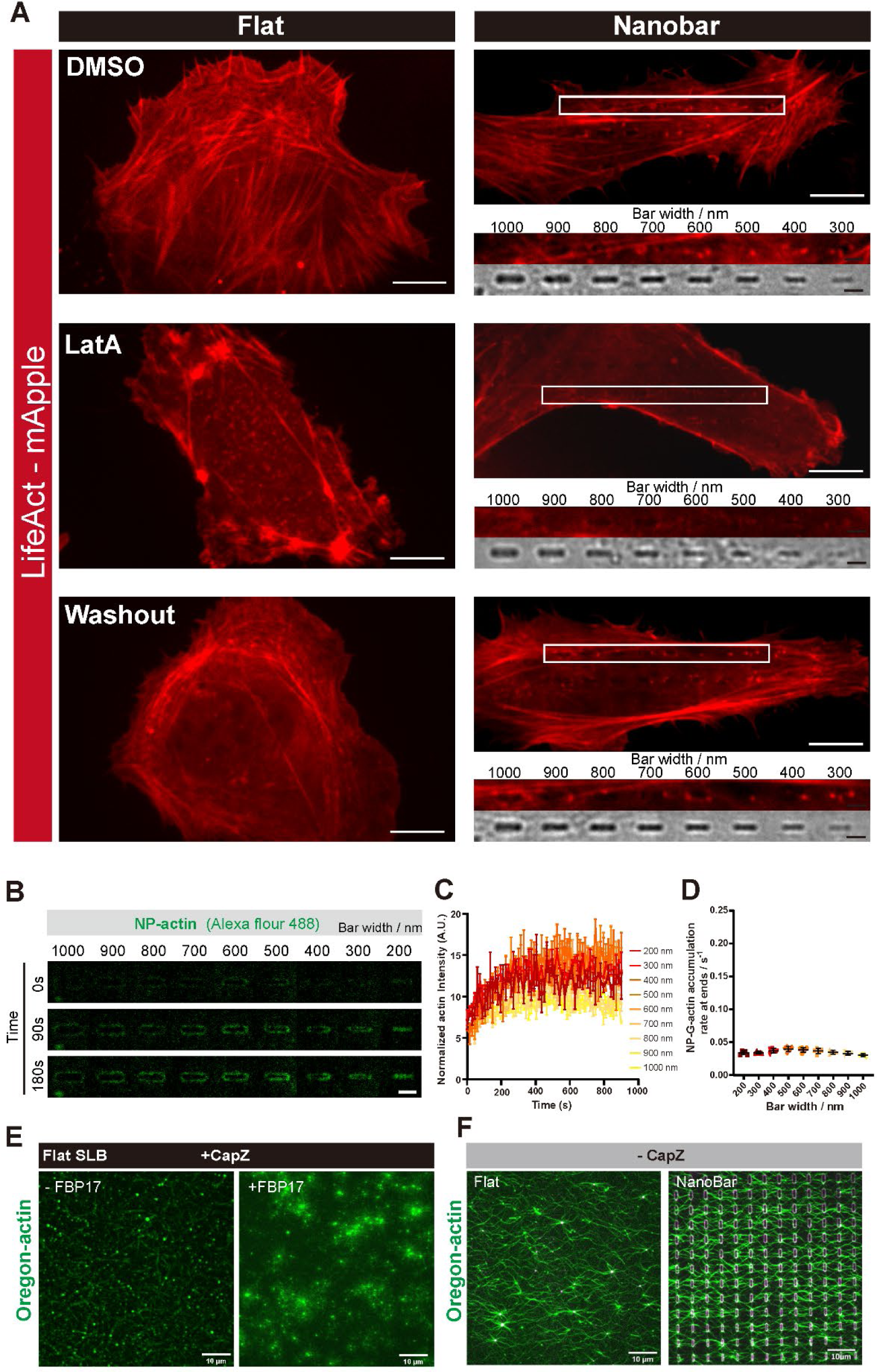
Dynamic radius-dependent actin assembly *in vivo* and *in vitro*. **A.** Confocal images of LifeAct-mApple-expressed U2OS live cells show F-actin recovery after 1h 100 nM LatA treatment and 20min after washout. An enrichment of repolymerized F- actin was observed on nanobars as compared with flat areas. Scale bar, 10 μm. Nano-bar panels: the 1.8X zoomed regions of nanobar array ranging from 300 to 1000nm in width (the white boxes). Scale bar, 2 μm. **B.** Averaged confocal images of *in vitro* reconstitution of 1.5 μM NP-actin (30% AF488 labeled) recruitment at 0/90/180s on the bilayer at nanobar of 200-1000 nm width with the presence of 200 nM FBP17, 50 nM N-WASP, and 5 nM Arp2/3. Scale bar, 2 μm. **C.** Normalized signal density of NP-actin based on their corresponding lipid bilayer intensity. Each point represents mean ± SEM from over 15 nanopillars. **D.** NP-G-actin accumulation rate at nanobars of 200-1000 nm width. (The linear fitted slope within 105-300s range in **C**). **E.** Confocal images of *in vitro* reconstitution of 1.5 μM actin (10% Oregon labeled) polymerization on the flat SLB with the presence of (-/+200 nM FBP17), 50 nM N-WASP, 12 nM CapZ and 5 nM Arp2/3. Scale bar, 10 μm. **F.** Confocal images of *in vitro* reconstitution of 1.5 μM actin (10% Oregon labeled) polymerization on the flat SLB (left) or patterned SLB(right) with the presence of 200 nM AF647-FBP17, 50 nM N-WASP, and 5 nM Arp2/3. Scale bar, 10 μm.

**Supplementary Figure 4.**
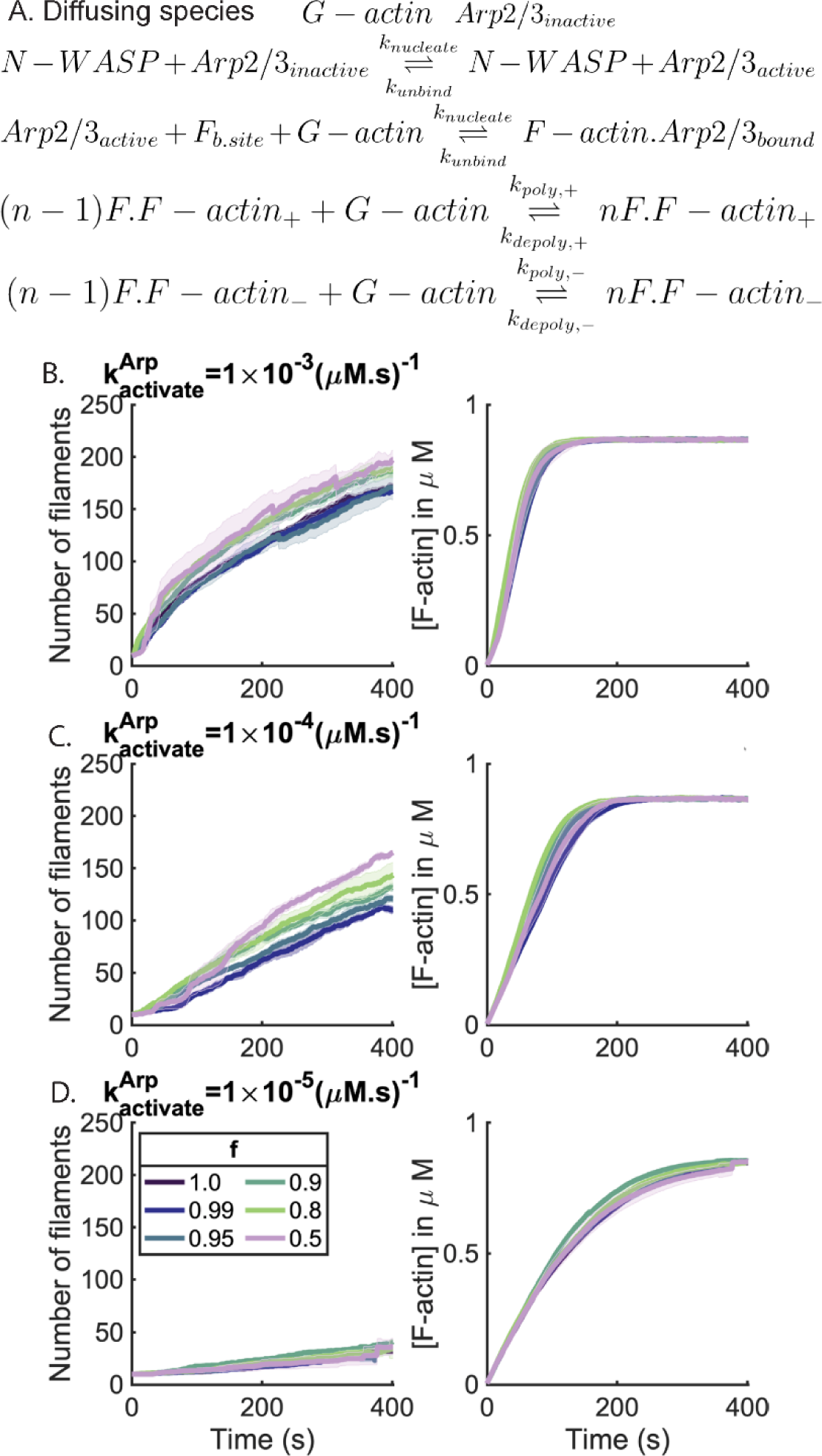
Spatial localization of N-WASP and N-WASP-driven activation rates are critical to enhance localized nucleation. A. The set of chemical reactions considered in MEDYAN are shown. G-actin and inactive Arp2/3 are allowed to diffuse throughout the reaction volume. Arp2/3 is activated proportional to the local N-WASP concentration. Active Arp2/3 can bind to a binding site of F-actin (one per 10 monomers in this study) to nucleate a new offspring filament. In addition, (de)polymerization reactions at both plus and minus ends of filaments are considered. B. -D) Activation rate of Arp2/3 plays a critical role in controlling the cooperative nucleation process. Plots show mean and standard error of mean plots at various values of localization factor (f) as time series. Each row corresponds to a particular value of activation rate.

**Supplementary Figure 5.**
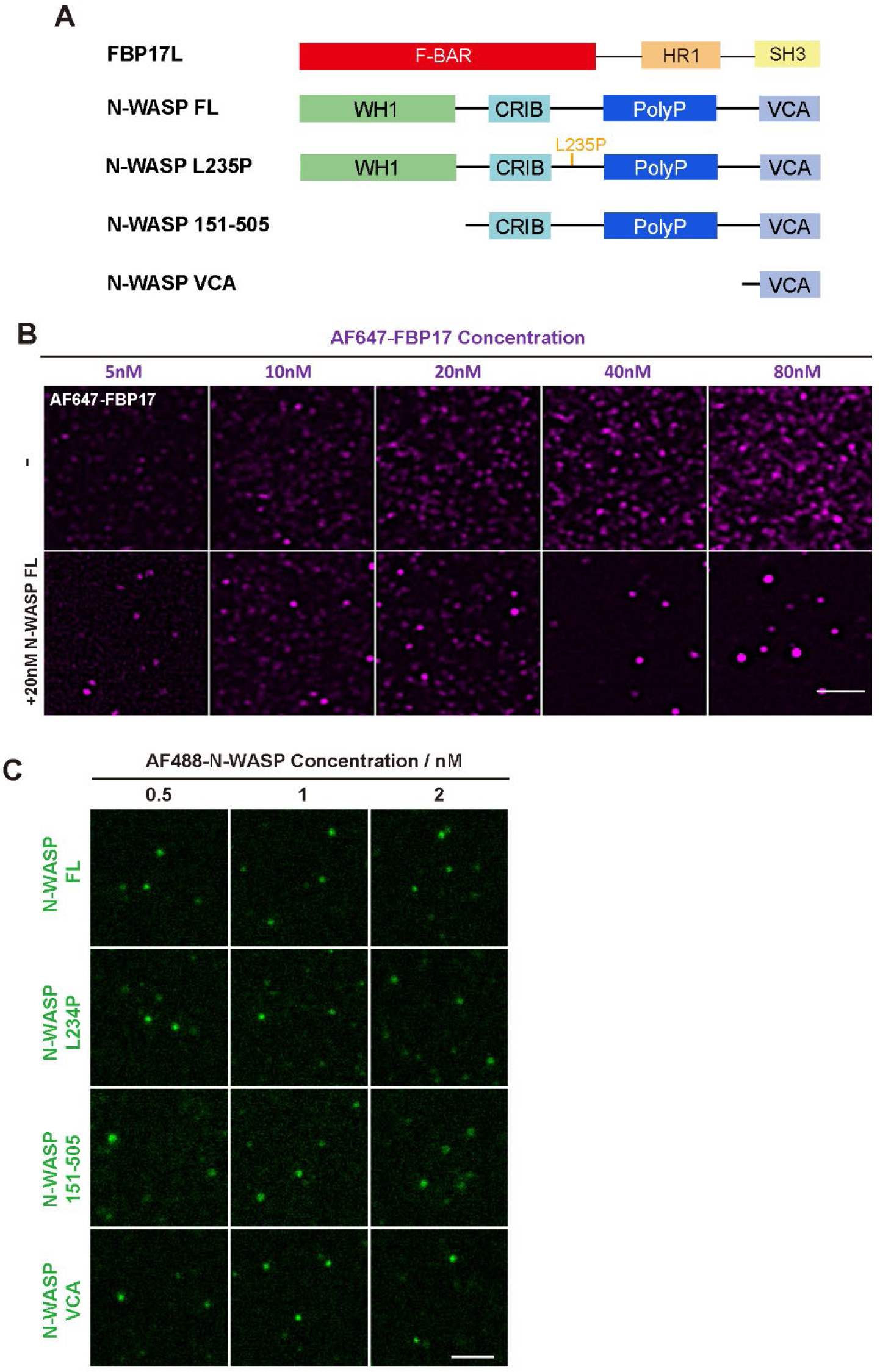
AF647-FBP17 and AF488-N-WASP single particle images. **A.** Domain illustration of FBP17 and N-WASP truncations. **B.** TIRFM single particle images of AF647-FBP17 at 5-80 nM on SLB with or without the presence of 20 nM FBP17. Scale bar, 5 μm. **C.** TIRFM single particle images of AF488-N-WASP truncation at 0.5, 1, and 2 nM on SLB. Scale bar, 2 μm.

**Supplementary Figure 6.**
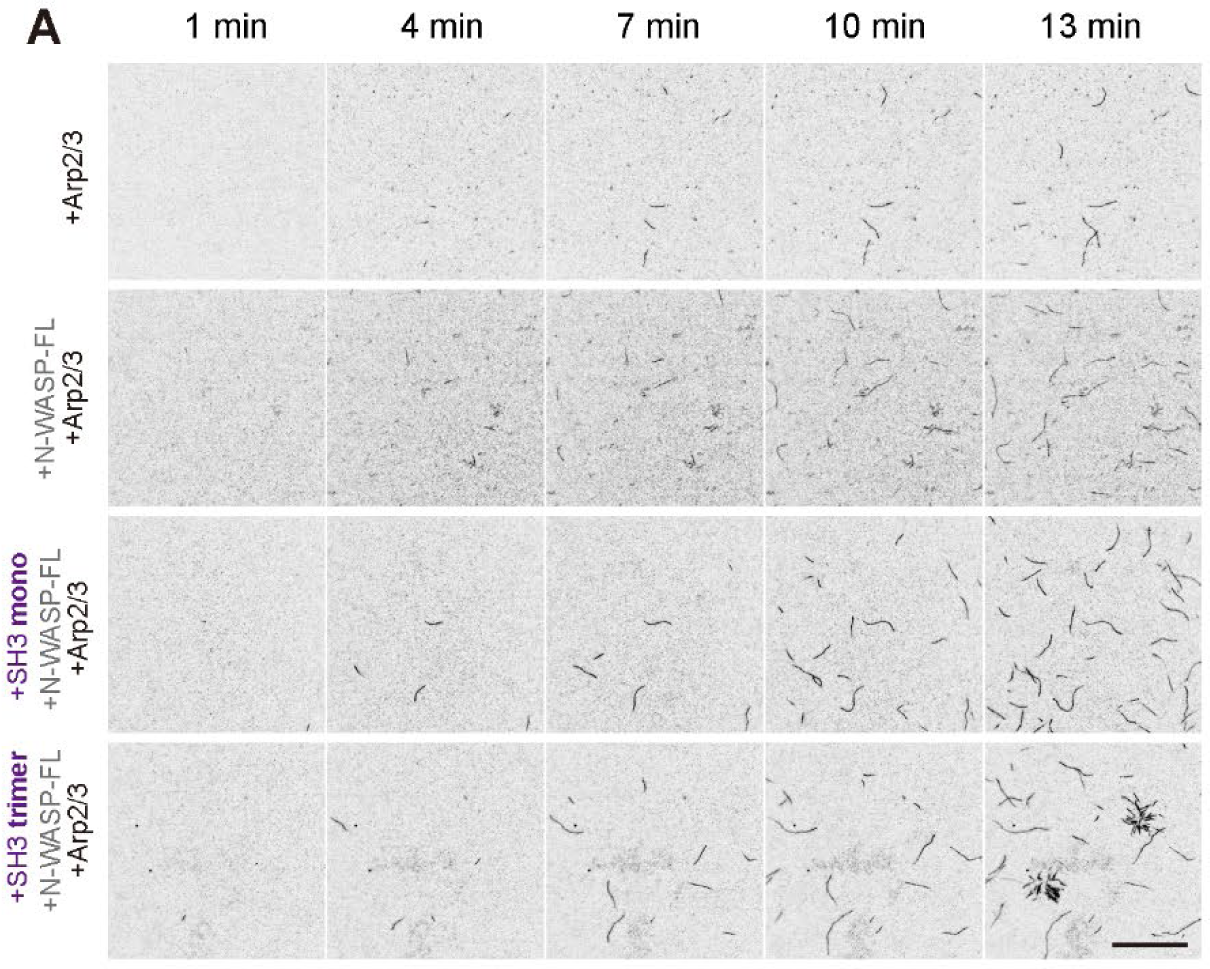
TIRF actin assembly assay of engineered SH3FBP17 A. TIRF images of actin polymerization (10% Oregon-labeled) with different combinations of 80 nM FBP17/oTri-SH3, 20 nM N-WASP FL and 5 nM Arp2/3. Scale bar, 10 μm.

**Table S1.**
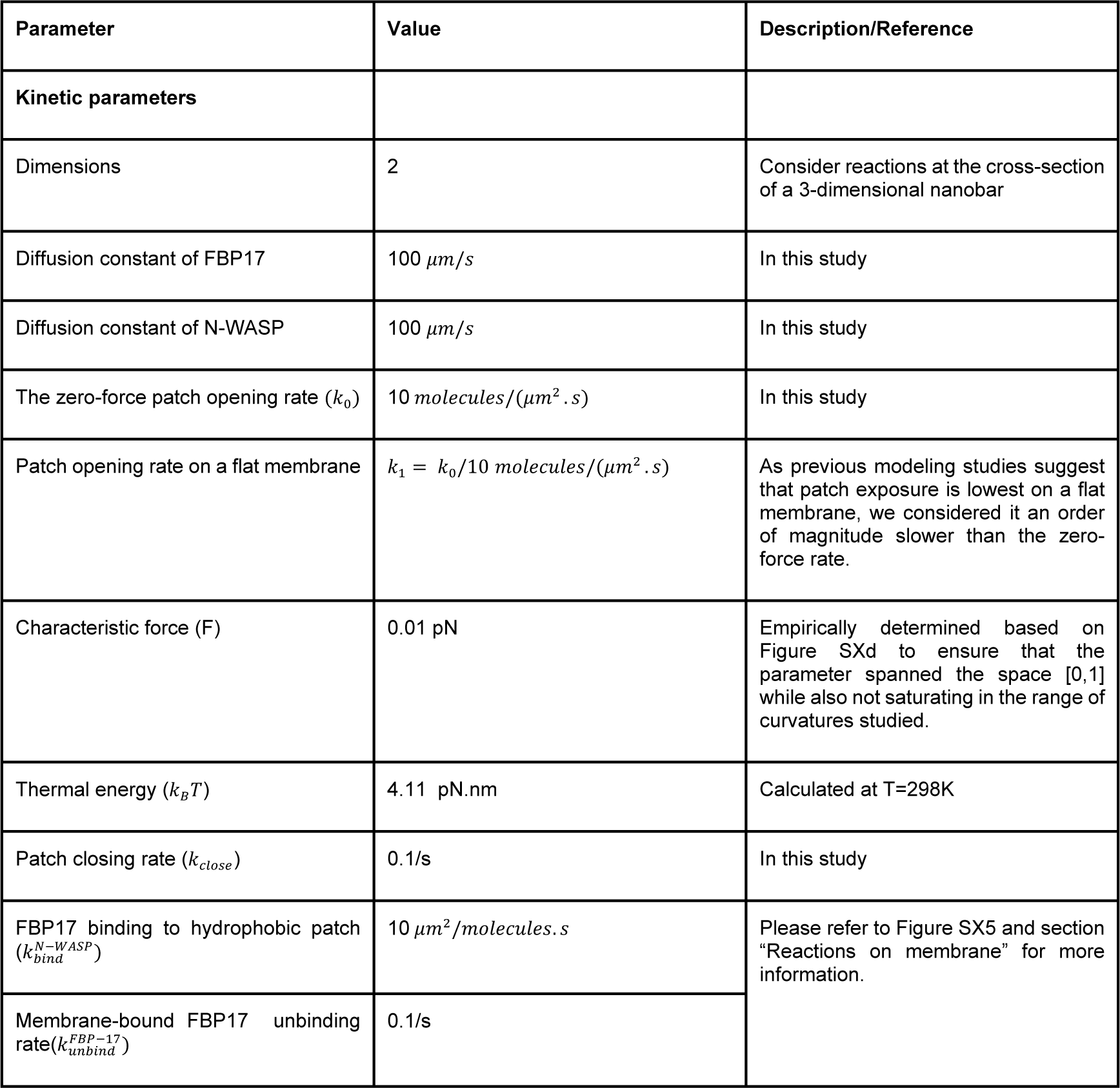
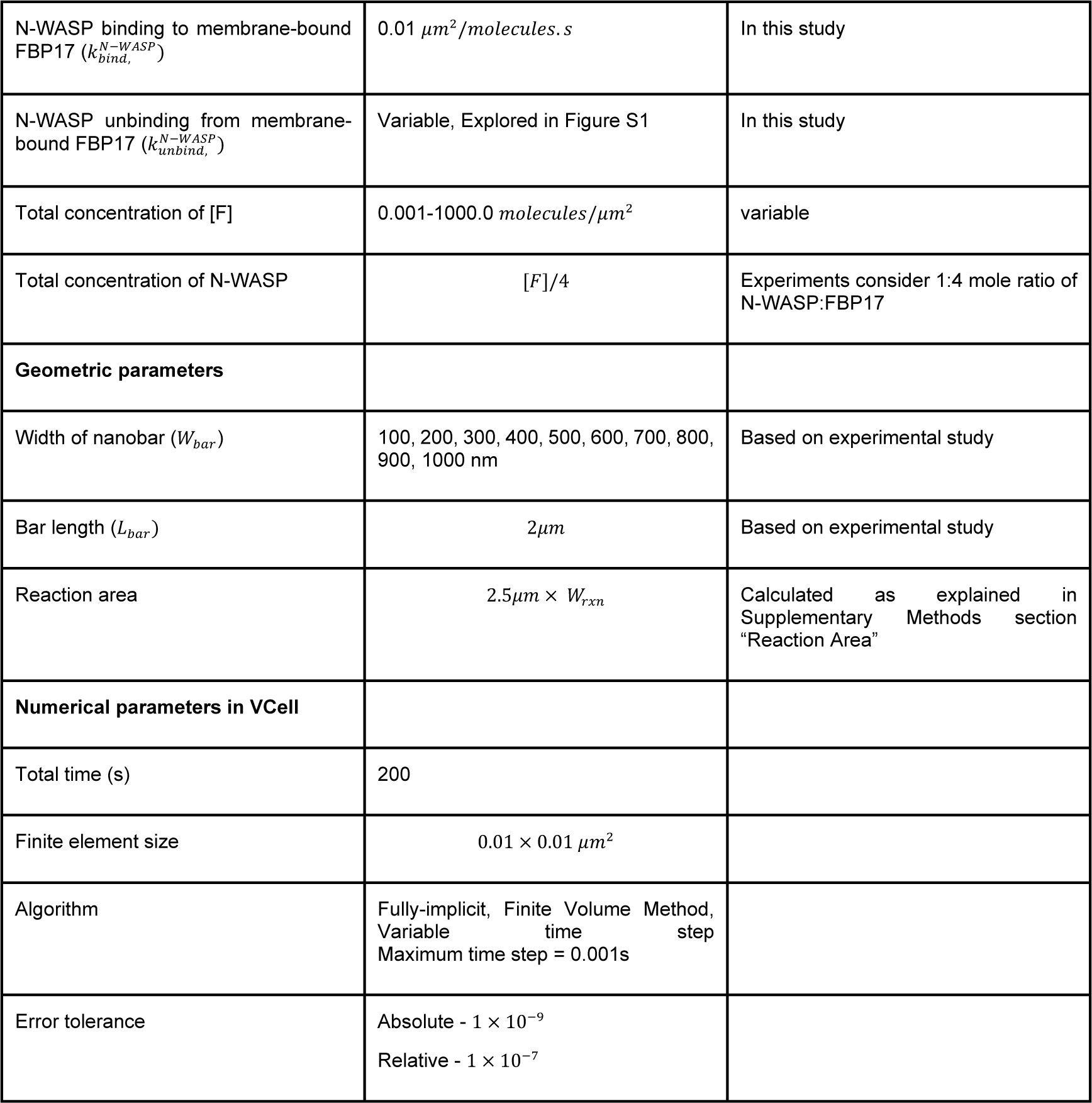
Table of parameters used to model curvature sorting of FBP17 and N-WASP on nanobars.

| Parameter | Value | Description/Reference |
| --- | --- | --- |
| **Kinetic parameters** |  |  |
| Dimensions | 2 | Consider reactions at the cross-section of a 3-dimensional nanobar |
| Diffusion constant of FBP17 | $100 \mu\text{m}/\text{s}$ | In this study |
| Diffusion constant of N-WASP | $100 \mu\text{m}/\text{s}$ | In this study |
| The zero-force patch opening rate ($k_0$) | $10 \text{ molecules}/(\mu\text{m}^2 \cdot \text{s})$ | In this study |
| Patch opening rate on a flat membrane | $k_1 = k_0/10 \text{ molecules}/(\mu\text{m}^2 \cdot \text{s})$ | As previous modeling studies suggest that patch exposure is lowest on a flat membrane, we considered it an order of magnitude slower than the zero-force rate. |
| Characteristic force (F) | 0.01 pN | Empirically determined based on Figure SXd to ensure that the parameter spanned the space $[0,1]$ while also not saturating in the range of curvatures studied. |
| Thermal energy ($k_B T$) | 4.11 pN.nm | Calculated at $T=298\text{K}$ |
| Patch closing rate ($k_{close}$) | 0.1/s | In this study |
| FBP17 binding to hydrophobic patch ($k_{bind}^{N-WASP}$) | $10 \mu\text{m}^2/\text{molecules} \cdot \text{s}$ | Please refer to Figure SX5 and section "Reactions on membrane" for more information. |
| Membrane-bound FBP17 unbinding rate ($k_{unbind}^{FBP-17}$) | 0.1/s | |
| N-WASP binding to membrane-bound FBP17 ($k_{bind}^{N-WASP}$) | $0.01 \mu m^2 / molecules.s$ | In this study |
| N-WASP unbinding from membrane-bound FBP17 ($k_{unbind}^{N-WASP}$) | Variable, Explored in Figure S1 | In this study |
| Total concentration of [F] | $0.001-1000.0 molecules/\mu m^2$ | variable |
| Total concentration of N-WASP | $[F]/4$ | Experiments consider 1:4 mole ratio of N-WASP:FBP17 |
| **Geometric parameters** |  |  |
| Width of nanobar ($W_{bar}$) | 100, 200, 300, 400, 500, 600, 700, 800, 900, 1000 nm | Based on experimental study |
| Bar length ($L_{bar}$) | $2\mu m$ | Based on experimental study |
| Reaction area | $2.5\mu m \times W_{rxn}$ | Calculated as explained in Supplementary Methods section "Reaction Area" |
| **Numerical parameters in VCell** |  |  |
| Total time (s) | 200 |  |
| Finite element size | $0.01 \times 0.01 \mu m^2$ | |
| Algorithm | Fully-implicit, Finite Volume Method, Variable time step<br>Maximum time step = 0.001s |  |
| Error tolerance | Absolute - $1 \times 10^{-9}$<br>Relative - $1 \times 10^{-7}$ | |

**Table S2.** Table of simulation parameters used in MEDYAN to simulate dendritic actin network. The mesoscopic rates used are shown within square brackets. aWe assume N-WASP to be membrane-bound and assume that the N-WASP does not diffuse during the timescale of the simulation.

| Parameter | Value | Comment/Reference |
| --- | --- | --- |
| **Geometric parameters** |  |  |
| Compartment size ($L_{comp}$) | 500 nm | 49 |
| Number of compartments in each dimension ($N_x \times N_y \times N_z$) | $16 \times 16 \times 1$ | - |
| Length of filament segment cylinder $L_{cyl}$ | 27nm (10 subunits) | - |
| Binding sites per cylinder $N_{b,site}$ | 1 | - |
| **Diffusion rates** |  |  |
| G-actin | $D_{actin} = 20 \mu m^2/s$<br>$k_{diff,actin} = 80 /s$ | 49 |
| N-WASP | $D_{NWASP} = 0 \mu m^2/s$ | a |
| Arp2/3 | $k_{diff,Arp2/3}$<br>$= k_{diff,actin}/10$ | 83,84 |
| **Kinetic rate constants** |  |  |
| Actin polymerization at plus ends ($k_{poly,+}$) | $11.6 (\mu M, s)^{-1} [0.154 s^{-1}]$ | 85 |
| Actin depolymerization at plus ends ($k_{depoly,+}$) | $1.4 s^{-1}$ | 85 |
| Actin polymerization at minus ends ($k_{poly,-}$) | $1.3 (\mu M, s)^{-1} [0.017 s^{-1}]$ | 85 |
| Actin polymerization at minus ends ($k_{depoly,-}$) | $0.8 s^{-1}$ | 85 |
| Arp2/3 activation rate ($k_{Arp2/3,activate}$) | 0.075, 0.0075, 0.00075<br>$(\mu M, s)^{-1} [10^{-3}, 10^{-4}, 10^{-5}] s^{-1}$ | In this study |
| Arp2/3 deactivation rate ($k_{Arp2/3,deactivate}$) | $0.01 s^{-1}$ | In this study |
| Arp2/3 binding rate ($k_{Arp2/3,bind}$) | $10.0 (\mu M^2, s)^{-1} [0.0017 /s^{-1}]$ | ODE model in <sup>83</sup> |
| Arp2/3 unbinding rate ($k_{Arp2/3,unbind}$) | $0.02 s^{-1}$ | 86 |
| **Concentrations** |  |  |
| Total Actin | $1 \mu M$ (19273 molecules) | - |
| Total N-WASP | $1 \mu M$ (19273 molecules) | - |
| Total Arp2/3 | 10 nM (198 molecules) |  |
| **Mechanochemical constants** |  |  |
| Arp2/3 unbinding force | $F_{branch,unbind} = 6pN$ | 87 |
| Characteristic force of Brownian ratchet | $F_{0,ratchet} = 1.5 pN$ | 88 |
| **Mechanical constants** |  |  |
| Actin filament stretching constant ($k_{str}$), and rest length $l_0$ | $100pN/nm$<br>$l_0 = L_{cyl}$ | 49 |
| Actin filament bending energy ($k_{bend}$) | $2690pN.nm$ | 89 |
| Boundary repulsion energy ($\epsilon_{boundary}$) | $10k_B T$ | - |
| Boundary repulsion screening length ($\lambda$) | 2.7 nm | - |
| Arp2/3 stretching constant ($k_{branch,stretch}$), and rest length $L_0$ | $k_{branch,stretch} = 100 pN/nm$,<br>$L_0 = 6 nm$ | 84 |
| Arp2/3 bending constant, I ($k_{branch,bend,I}$), equilibrium angle | $10 pN.nm, \pi/2$ | 84 |
| Arp2/3 bending constant, II ($k_{branch,bend,II}$), equilibrium angle | $20 pN.nm, 70^\circ$ | 84 |
| Arp2/3 dihedral constant ($k_{branch,dihedral}$) | $10 pN.nm$ | 84 |
| **Minimization parameters** |  |  |
| Time scale of chemical steps | 5 ms |  |
| Force tolerance for mechanical minimization | 10 pN |  |
| Seed filaments |  |  |
| Number | 10 |  |
| Length | 27 nm |  |
| **Simulation parameters** |  |  |
| Total time | 400 s |  |
| Replicates | 3 |  |

**Table S3.**

| Plasmid No. | Name | Region | Source |
| --- | --- | --- | --- |
| pMYS1601 | pET-28a-His-FBP17 | 1-617aa of FBP17 | Min Wu Lab |
| pMYS1602 | pET-32m-N-WASP FL | 1-505aa of N-WASP (Homo sapiens) | Mingjie Zhang lab |
| pMYS1603 | pET-32m-N-WASP L235P | L235P point mutation of N-WASP FL | Mingjie Zhang lab |
| pMYS1604 | pET-32m-N-WASP 151-505 | 151-505aa of N-WASP | Mingjie Zhang lab |
| pMYS1605 | pET-32m-N-WASP VCA | 392-505aa of N-WASP | This paper |
| pMYS1606 | pNIC-Bsa4-His-avi-SH3 | SH3 domain of FBP17 | This paper |
| pMYS1607 | pNIC-Bsa4-His-avi-oTri-SH3 | SH3 domain of FBP17+Trimer CC sequence | This paper |

## Notes

### Competing Interest Statement

The authors have declared no competing interest.

